# Pollinator mosaics mirror floral trait divergence within and between species of *Castilleja*

**DOI:** 10.1101/2022.07.04.498476

**Authors:** Katherine E. Wenzell, Krissa A. Skogen, Jeremie B. Fant

## Abstract

1. Pollinator interactions are important in the evolution of floral traits, given that pollinators can directly influence plant fitness and mating patterns through interactions with flowers. However, geographic variation in both plant traits and floral visitors across multiple populations is largely understudied, despite being ubiquitous. This study explores whether a geographic mosaic of ecological interactions underlies observed patterns of floral divergence 1) among species of the *Castilleja purpurea* complex (*C. purpurea, C. citrina*, and *C. lindheimeri*) and the congener *C. sessiliflora*, as well as 2) within *C. sessiliflora*, across its wide geographic range. We sampled floral visitors and floral traits (morphology and color) at 23 populations across a 1900 km study area in 1-3 years, with reproductive fitness (fruit set) data for 18 of these populations.
2. We documented a wide diversity of pollinator functional groups visiting the four focal species, including bees, butterflies, hawkmoths, and hummingbirds. Visitor assemblages varied among species and across geography in the composition and diversity of floral visitors. We found relationships between floral traits and visitation by certain pollinator groups, which often aligned with syndrome-associated predictions. Additionally, we found evidence that visitation from pollinators predicted via syndromes was associated with increased reproductive fitness for two species: the red-flowered *C. lindheimeri* and hummingbird visitors, and the long-floral-tubed *C. sessiliflora* and hawkmoths.
3. Beyond these cases, we found that pollinator functional groups were not restricted to plant species, and instead pollinators were largely generalist in their foraging behavior, suggesting the likelihood for incomplete reproductive isolation and the potential for ongoing gene flow among plant species where their ranges overlap.
4. This study provides a large-scale exploration of how variation in pollinator assemblages across distributions may underlie floral trait divergence within and among recently diverged species, even when characterized by largely generalized modes of pollination. Our extensive sampling of 23 populations over multiple years across a large geographic area highlights the value of range-wide studies for characterizing patterns of divergence and speciation mediated by ecological interactions.

## Introduction

The role of animal pollinators in angiosperm speciation is well-studied (Johnson, 2006), from macroevolutionary (Crepet, 1984; Lunau, 2004; van der Niet and Johnson, 2012), microevolutionary (Gervasi and Schiestl, 2017) and ecological (Campbell et al., 1997; Fulton and Hodges, 1999; Bradshaw and Schemske, 2003) perspectives. However, discussion persists over the degree to which pollinator identity and shifts between pollinators may confer reproductive isolation, and thus may represent a mechanism to drive plant speciation and floral diversification (Waser and Campbell, 2004; Kay and Sargent, 2009; van der Niet et al., 2014). Strong reproductive isolation due to pollinator shifts has been observed in sympatry (Schemske and Bradshaw, 1999), but such cases may be rare (Waser et al., 1996; Ollerton et al., 2009) and many well-known examples focus on allopatrically-diverged species experiencing secondary contact at the edges of their ecological tolerances (Campbell et al., 1997; Fulton and Hodges, 1999; Bradshaw and Schemske, 2003). While informative, these examples may reveal how reproductive isolation is maintained, but not necessarily how it first arises. Studies that characterize geographic variation in floral traits and pollinators both within and among recently diverged taxa are needed to better understand early stages of floral phenotype divergence and reproductive isolation, and the role of pollinator shifts due to geographic mosaics therein.

Models of pollinator-mediated plant speciation often invoke how geographic variation relates to processes of divergence, a key factor for studies of adaptation to geographically varying pollinator assemblages (Grant, 1949; Stebbins, 1970; Johnson, 2006). Prior work suggests that the locally most effective pollinator (Stebbins, 1970) will exert selection on floral traits, based on its morphology, physiology, and foraging behavior (Schiestl and Johnson, 2013), but that the most effective pollinator may vary across species distributions (Thompson, 2005), along with its ecological context and fitness contributions (Kay and Sargent, 2009; van der Niet et al., 2014; Ohashi et al., 2021). This variation may lead to divergence in reproductive traits and can give rise to reproductive isolation and speciation between populations. While divergence is most likely to occur in concert with other sources of ecological variation, especially at large geographic scales (Johnson, 2006; Nosil, 2012), pollinators are commonly expected to shape both trait divergence and reproductive isolation, given their direct influence on both selection (via reproductive fitness) and gene flow (via pollen movement) of flowering plants (Waser and Campbell, 2004).

While variability in both biotic and abiotic environments is well-documented across species distributions (Thompson, 2005), range-wide geographic variation in pollinator-mediated selection of floral traits is relatively understudied (Herrera et al., 2006), as such studies tend to focus on one or a few populations in hybrid zones or zones of contact (Bradshaw and Schemske, 2003; Campbell and Aldridge, 2006; Hopkins and Rausher, 2012). Despite an increasing focus on geographic mosaics of plant-pollinator interactions (Anderson and Johnson, 2008; Boberg et al., 2014; Ellis et al., 2021; Johnson et al., 2021), an integrated understanding of the impacts on speciation remains an important knowledge gap. In particular, characterizing the role of pollinators in mediating reproductive isolation at varying geographic scales (i.e., in sympatry, parapatry, or allopatry) requires studies of pollinator mosaics at such scales (Waser and Campbell, 2004), including within and among species across their distributions (Kay and Sargent, 2009). Of particular importance are range-wide studies of variation in local pollinators among recently diverged species or pollination ecotypes, which may represent recent or incipient speciation (Pellmyr, 1986; Oyama et al., 2010; van der Niet et al., 2014; Sobel and Streisfeld, 2015).

Here, it is useful to consider two early stages of speciation: phenotypic divergence and reproductive isolation. While there are cases of speciation that exhibit both stages (Coyne and Orr, 2004), increasing evidence suggests that, if selection is strong enough, phenotypic divergence can arise rapidly and may persist in the face of ongoing gene flow, especially in cases of ecological speciation (Nosil, 2008; Tavares et al., 2018). Given that genomic incompatibilities and/or hybrid inviability (underlying strong reproductive isolation) are expected to take considerable time to arise in isolation, early or intermediate stages of ecological speciation may be more likely characterized by phenotypic divergence, driven by strong, recent selection, with weak reproductive isolation and ongoing gene flow. Phenotypic and genetic patterns consistent with this scenario have been reported in systems with phenotypic divergence in reproductive traits despite a lack of genetic structure or differentiation (Nosil and Crespi, 2004; Mason and Taylor, 2015; Harris et al., 2018) and include examples of pollinator-mediated selection on floral traits (Streisfeld and Kohn, 2005; Whibley et al., 2006; Hopkins et al., 2012; Stankowski et al., 2017).

Recent work on the species of the *Castilleja purpurea* (Nutt.) G. Don species complex, and its congener *C. sessiliflora* Pursh demonstrated high levels of floral trait variation despite low genetic differentiation within and among these species (Wenzell et al., 2021). *Castilleja sessiliflora* displays geographic variation in floral traits across its wide range, with much of this variation concentrated in inflorescence color: white-green to pale pink inflorescences from north to south, with distinct bright pink and yellow morphs in the southern range extent (Fig. 1). In contrast, the three species of the *C. purpurea* complex, recently elevated to species status, have relatively small, overlapping ranges and vary primarily in inflorescence color: *C. purpurea* has purple bracts, *C. citrina* Pennell has yellow bracts, and *C. lindheimeri* A. Gray has red-orange bracts (Nesom and Egger, 2014). Despite these striking differences in color, the species are characterized by low levels of differentiation at targeted genomic loci, even across narrow clines in floral color (Wenzell et al., 2021), suggesting these species are recently diverged, possibly due to selection on floral color that could be mediated by pollinators.

**Figure 1.**
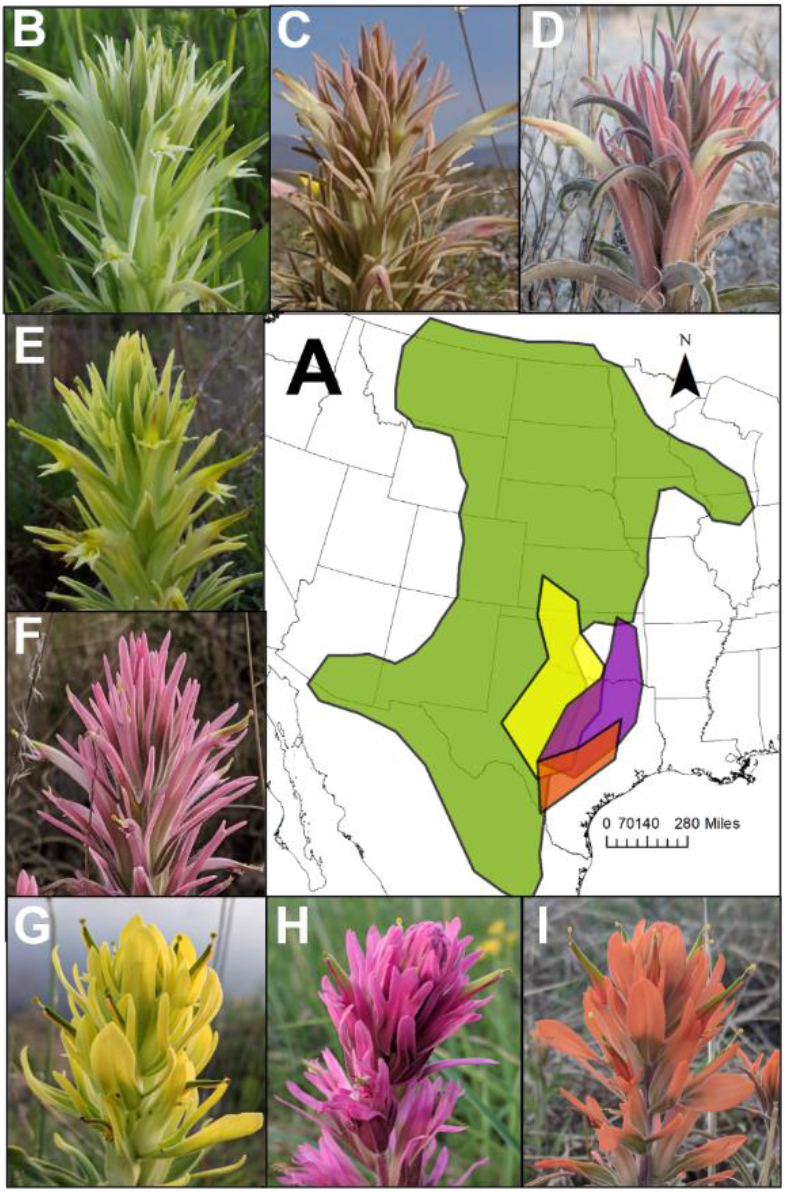
Geographic range (A) and floral trait divergence within *Castilleja sessiliflora* (B-F; green range on map) and among species of the *C. purpurea* complex: *C. citrina* (G, yellow range), *C. purpurea* (H, purple range), and *C. lindheimeri* (I, orange range). Floral trait variation within *C. sessiliflora*: a grade of white-green inflorescences in the northeastern range extent (B) to pale pink in the southwest (C, D), and two distinct floral morphs in the southern range, with shorter corollas and bright yellow (E, population SMP) or pink (F, SIC) inflorescence color (Wenzell et al. 2021). Photos by K. Wenzell.

In this study, we characterize patterns of pollinator visitation and diversity range-wide to investigate whether floral divergence may be driven by selection mediated by pollinators. We address the research question: Does a mosaic of pollinator visitation underlie the patterns of divergence seen in floral traits for *C. sessiliflora* and the *C. purpurea* complex? We assess this question both for among-species floral divergence (i.e., among all four study species: *C. purpurea, C. citrina, C. lindheimeri*, and *C. sessiliflora*), as well as for within-species floral divergence, focusing on variation across the wide range of *C. sessiliflora*. If a pollinator mosaic is driving floral divergence, we expect to find evidence for the following hypotheses: 1) pollinator visitation and diversity vary among species and across geography, 2) visitation from different pollinator functional groups is associated with variation in floral traits likely to be important for pollination, and 3) plant fitness will be higher with increased visitation from pollinator groups corresponding to floral traits. To test these hypotheses, we collected data on the composition of floral visitors to 23 natural populations across 1–3 years, in combination with data on floral traits and female fitness (fruit set) across the range of each of the four species. This study unites components of pollinator-mediated selection on floral traits to inform our understanding of how these factors interact across wide geographic scales, thus shaping floral divergence, a first step in pollinator-mediated speciation.

## Materials and Methods

### Study system

The genus *Castilleja* (paintbrushes, Orobanchaceae) is hemiparasitic and known for variability in color and morphology of flowers and showy floral bracts, which is often attributed to putative hybridization or the retention of ancestral polymorphisms following rapid radiation (Tank and Olmstead, 2008). This study focuses on four perennial species characterized by diverse floral traits: the widespread *C. sessiliflora* and the species of the more geographically restricted *C. purpurea* complex: *C. purpurea, C. citrina*, and *C. lindheimeri* (Fig. 1). Given their geographic proximity and morphological similarity, *C. sessiliflora* and the *C. purpurea* complex are expected to be close relatives (D. Tank pers. comm.). These species are largely self-incompatible, and flowers remain open for several days and offer both pollen and nectar rewards (K. Wenzell, unpublished data).

### Data Collection

#### Floral traits

Floral traits were measured at 23 focal populations and included 5 morphological traits (corolla length, corolla width, petaloid lip length, stigma exsertion, and bract lobe width) and inflorescence color (Royal Horticultural Society (RHS) floral color charts). Traits were measured from two flowers each of 30 plants in a single year at each population and are described in greater detail in Wenzell, et al. (2021). Because RHS color charts do not capture human-invisible UV reflectance, which is important for vision in many pollinating insects (Peitsch et al., 1992), we photographed inflorescences of each species and floral morph using an ultraviolet-sensitive camera (Canon EOS REBEL T3i camera with UV-transmitting lens and filter) under natural daylight conditions at 6 populations in 2017 (one each per species of the *C. purpurea* complex and three populations of *C. sessiliflora*, including both populations with distinct yellow and pink floral morphs).

To quantify floral color, RHS color codes were converted to Red-Green-Blue (RGB) values, followed by nonmetric multidimensional scaling (NMDS) of RGB values, using Gower distances in R package vegan (Oksanen et al., 2019), to make color values more easily interpretable. The resulting NMDS axes were associated with each component of RGB color using the envfit function and represent variation in inflorescence color throughout the study (Supplemental Fig. S1). Overall, the NMDS1 axis was characterized by warm colors (red, orange, yellow) at low values and cool colors (purple, pink) at high values. NMDS2 was roughly characterized by darker, reddish colors (purple, red) at high values and paler, greenish colors (green, yellow-green, yellow) at low values.

#### Sampling of pollinator observations

Pollinator observations were conducted for 1–3 years from 2017–2019 at 23 populations distributed across the range of each species (12 populations of *C. sessiliflora* and 11 populations representing the *C. purpurea* species complex: four populations of *C. purpurea*, four populations of *C. citrina*, and three populations of *C. lindheimeri*; Supplemental Table S1). Due to the challenge of sampling such a large geographic area, pollinator observations were conducted over one 24-hour period per population per year, and not every population could be sampled in each year. Specifically, three *C. sessiliflora* populations in the center of the range (SDC, SPB, and SRS) were sampled only in 2017, and one population of each of the *C. purpurea* complex species (CHCL, LMN, and PTMS) were sampled only in 2019. Furthermore, *C. sessiliflora* was the only species sampled in 2017 (Table 1). Complete information on sampling years and datasets for each population are given in Table S1.

**Table 1.** Floral visitation summary table. Number of recorded floral visits to each plant species for each pollinator functional group. For each plant species, the number of sampled populations, years sampled, and the number of observation datapoints (population-year-dataset observations, total N = 61) is given, along with total number of visits and average number of visits per datapoint (calculated as Total visits / datapoints per species).

|  | Small<br>/ Med<br>Bee | Large<br>Bee | Hawkmoth | Hummingbird | Butterfly | Other | Total<br>visits | Average<br>visits per<br>datapoint |
| --- | --- | --- | --- | --- | --- | --- | --- | --- |
| *C. citrina* | 85 | 268 | 49 | 63 | 53 | 1 | 519 | 47.2 |
| populations = 4 |  |  |  |  |  |  |  |  |
| years: 2018, 2019 |  |  |  |  |  |  |  |  |
| datapoints = 11 |  |  |  |  |  |  |  |  |
| *C. lindheimeri* | 4 | 0 | 20 | 430 | 115 | 2 | 571 | 71.4 |
| populations = 3 |  |  |  |  |  |  |  |  |
| years: 2018, 2019 |  |  |  |  |  |  |  |  |
| datapoints = 8 |  |  |  |  |  |  |  |  |
| *C. purpurea* | 248 | 85 | 409 | 144 | 124 | 42 | 1052 | 95.6 |
| populations = 4 |  |  |  |  |  |  |  |  |
| years: 2018, 2019 |  |  |  |  |  |  |  |  |
| datapoints = 11 |  |  |  |  |  |  |  |  |
| *C. sessiliflora* | 698 | 317 | 683 | 0 | 105 | 67 | 1870 | 60.3 |
| populations= 12 |  |  |  |  |  |  |  |  |
| years: 2017, |  |  |  |  |  |  |  |  |
| 2018, 2019 |  |  |  |  |  |  |  |  |
| datapoints = 31 |  |  |  |  |  |  |  |  |

At each population, two observers recorded floral visits simultaneously during eight 20-minute observation periods during daylight hours (evenly spaced throughout periods of pollinator activity) and one 60-minute observation period at dusk, which started 15 minutes before sunset. This sampling resulted in 440 observer minutes recorded per site per year, for a total of 330 hours of observation time across all sites and years. Floral visitors were recorded to pollinator functional group (Fenster et al., 2004) and were analyzed as such, though genus (e.g., *Bombus* sp.) or species (e.g., *Hyles lineata*) was noted when possible. Pollinator functional groups included: hawkmoths (Sphingidae), hummingbirds (Trochilidae), bumblebees (*Bombus* sp.), other bees classified as large, medium, or small, bee flies (Bombyliidae), non-Sphingid Lepidoptera categorized as small or large butterflies or other moths, and flies (Diptera; additional information can be found in Supplemental Methods). For analysis, infrequent groups (those recorded visiting < 50 flowers in the total dataset or recorded at only one population) were pooled with more common, functionally similar groups or into a group of “other” visitors. This latter category includes mainly bee flies, flies, and non-Sphingid moths. Final analyses included the following pollinator functional groups: hawkmoths, hummingbirds, bumblebees and large bees, small and medium bees, butterflies, and other visitors.

In all years (2017–2019), floral visitation data were recorded in the following way: a focal patch of flowering *Castilleja* plants was designated within approximately 1–2 m of the observer (near enough to observe and record visitation by sweat bees or other small insects), and visitation to flowers of these focal plants was recorded during all observation periods, which represents the “narrow-view dataset.” The number of open flowers on each focal plant was recorded for each day of observations. However, it was noted that certain pollinator functional groups (especially hummingbirds) were wary of approaching plants close to observers, which had the effect of excluding them from this dataset. To account for this, an additional, second visitation dataset was collected in 2019. For this complementary “wide-view dataset”, observers recorded floral visits to any *Castilleja* plants in their field of view that occurred during observation periods. The approximate number of flowering stems of *Castilleja* in this wide field of view was counted to generate an estimate of the number of open flowers available to pollinators (calculated as the average number of open flowers per focal plant, multiplied by the number of flowering stems in the wide view). Visits to narrow-view focal plants were not included in an observers’ wide-view dataset, and care was taken to avoid observers sharing the same wide view, to prevent double-counting of floral visits. Unless otherwise noted, floral visitation data from both observation methods (referred to as “dataset types” hereafter) are included in analyses, and dataset type was included as a fixed effect in our models to account for the potential impact of these different observation methods.

For both methods, observers recorded the number of flowers visited by a given pollinator during an observation period when a flower was visibly probed and the visitor appeared to contact the anthers and/or stigma. Number of floral visits by each pollinator functional group was pooled across observation periods for a given population-year, and observation periods were an equal amount of time for each population-year. Data were calculated as count (number of floral visits by pollinator group per population-year), visitation rate (number of floral visits relative to the total number of open flowers available in the narrow or wide view, depending on dataset, per hour), and as proportion of all visits (number of floral visits by one pollinator group relative to the total number of visits from all pollinator groups per population-year; Supplemental Table S2). Because sampling time (observer hours) was equal across population-years, and to compare relative contribution of different pollinator functional groups across populations, subsequent analyses use proportion data unless otherwise noted. Calculations and analyses were performed in R (R Development Core Team, 2019) using tidyverse packages (Wickham et al., 2019).

#### Plant fitness measurements

To characterize the reproductive fitness of focal populations, we measured fruit to flower ratio in a subset of 18 populations (Table S1), based on approximately 30 plants per population. Some populations were sampled in multiple years for a total of 20 population-year observations and a total sample size of 591 plants. Fruit set was measured as the total number of filled fruits (or enlarging ovaries if populations were sampled prior to fruit maturation) divided by the number of total flower nodes (1 flower/node) along one or more flowering stems per individual, *i*.*e*. the proportion of filled fruits per individual. This fruit set value was used as an estimate of plant fitness.

### Variation in key floral traits among species and populations

To establish overall variation of floral traits before proceeding with tests of individual traits, we performed a MANOVA with all seven measured traits using the manova() function in R, testing for multivariate variation among species and among populations nested within species, with individual plant as the sampling unit (N = 684). Next, we identified which individual floral traits varied among species and among populations (nested within species) by comparing mean values of each trait using ANOVA (aov() function). For traits that varied significantly among species, we then used post hoc Tukey HSD tests (TukeyHSD() with 95% CI) to make pairwise comparisons among species.

### How does pollinator visitation vary among plant species and across geography?

We first assessed how the composition of floral visitors varied among species. We tested whether each pollinator functional group varied significantly in its number and proportion of total visits to each plant species using generalized linear mixed models (GLMMs) in R package glmmTMB (Magnusson et al., 2020), followed by Type II Wald Chi-Square tests of the model output, using function Anova() in package car (Fox et al., 2020). Multiple comparison analyses were then performed with function lsmeans() in package lsmeans, using pairwise comparisons among plant species with Tukey adjustments (Lenth, 2018). Separate models were run for each pollinator functional group, and models were run using number or proportion of visits, which were calculated as individual datapoints for each population-year sampled and each dataset type (N = 61), with dataset type included as a fixed effect and observation year as a random effect. To assess whether count and proportion data revealed different patterns, both measures were analyzed using separate models; methods for count models are detailed in Supplemental Methods. For proportion models, proportion of visits for each pollinator type was the response, weighted by total number of visits, with plant species as the predictor, and a betabinomial error distribution. To assess potential overdispersion in the data, we ran models using both a binomial and betabinomial error distribution and used package DHARMa (Hartig, 2016) to examine residuals and assess evidence for overdispersion in each model. The models with binomial distributions consistently showed evidence of overdispersion, while the models using betabinomial distributions did not, and thus the latter were chosen for models of proportion data throughout the study unless otherwise noted.

For each population, we visualized the overall proportion of visits across years, calculated as the total number of visits from each pollinator group, relative to the total number of visits (following Crawley, 2015 p. 257; Fig. 2). We performed statistical analyses of geographic variation in pollinator visitation only in the widespread *C. sessiliflora*, given the more restricted geographic ranges and limited number of sampled populations for the species of the *C. purpurea* complex. For these tests, we performed GLMMs as described above with proportion of visits of each pollinator type as the response term and population latitude as predictor, along with dataset type as a fixed effect and year as a random effect. Latitude was used as a proxy for geographic variation across the range of *C. sessiliflora*, given the large latitudinal spread of sampled populations.

**Figure 2.**
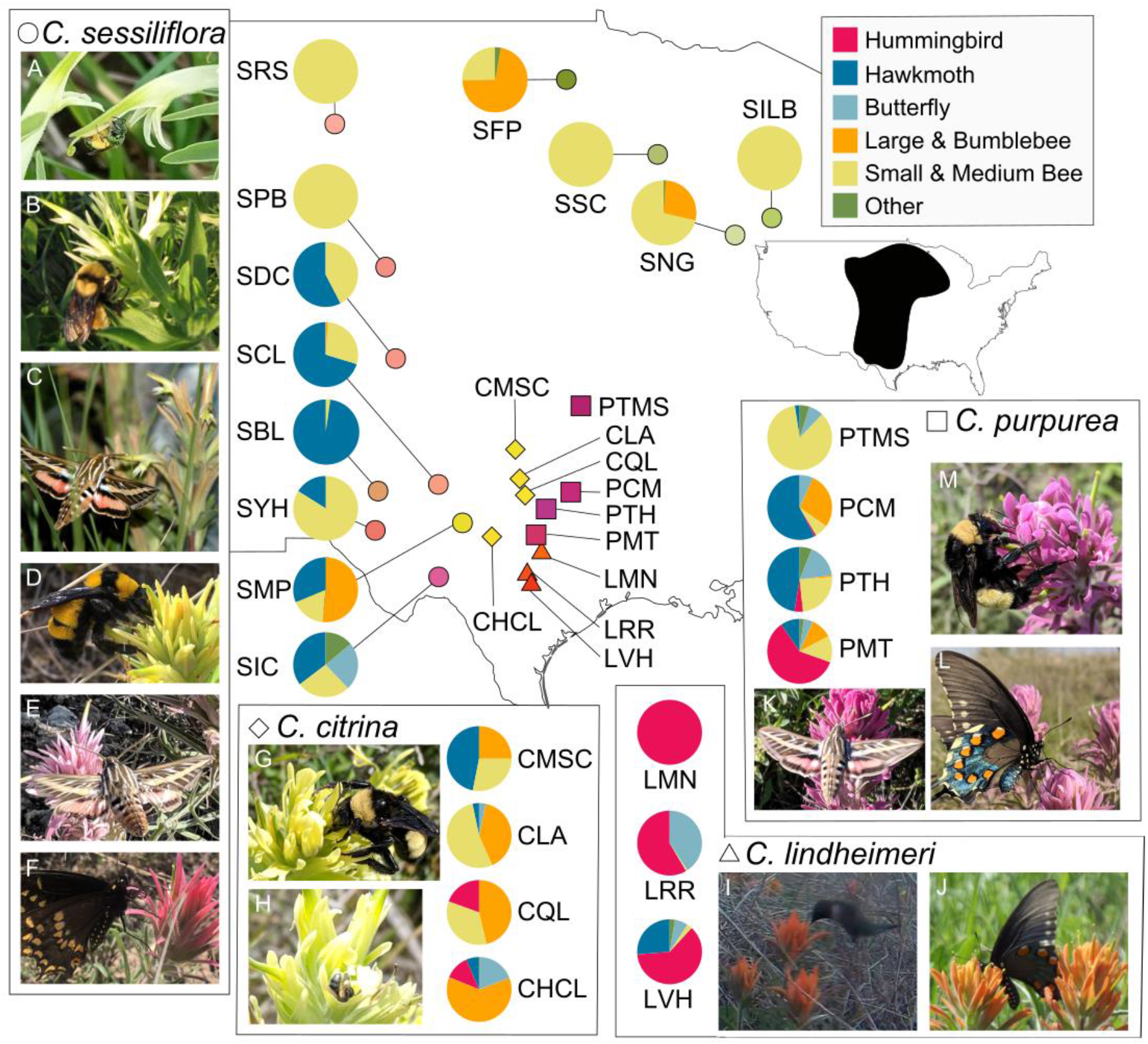
Geographic mosaic of floral color and floral visitors across the ranges of *C. sessiliflora* and the *C. purpurea* complex. Shapes on map show 23 sampled populations of *C. sessiliflora* (circles), *C. purpurea* (squares), *C. citrina* (diamonds), and *C. lindheimeri* (triangles); fill color is population median floral color (median RGB values; N= 684). Pie charts show overall proportion of visits by pollinator functional group to populations across years. Photos show selected floral visitors: A) small sweat bee (Halictidae) collecting pollen on *C. sessiliflora* at SNG; B) bumblebee (*Bombus fervidus*) nectaring at SNG (photo: J. Fant); C) hawkmoth (*Hyles lineata*) in Colorado, near SDC (photo: S. Todd); D) bumblebee (*Bombus sonorus*) on the yellow floral morph at SMP (photo: S. Deans); E) hawkmoth (*H. lineata*) and F) butterfly (black swallowtail, *Papilio polyxenes*) on the pink floral morph at SIC (photo: S. Deans); G) bumblebee (*B. pensylvanicus*) and H) small bee on *C. citrina* at CQL; I) hummingbird at LRR and J) pipevine swallowtail butterfly (*Battus philenor*) on *C. lindheimeri*; K) hawkmoth (*H. lineata*) visiting *C. purpurea* at PTH; L) pipevine swallowtail butterfly (*B. philenor*) and M) bumblebee (*Bombus pensylvanicus*) at PCM. Photos by K. Wenzell unless otherwise noted.

#### Pollinator Diversity Index

Due to the difficulty in comparing categorical variation in pollinator assemblages across species (i.e., the presence of absence of different pollinator functional groups), we summarized variation in overall diversity of pollinator assemblages by calculating a Pollinator Diversity Index in observed visitation by functional group (adapted from Lázaro et al., 2009). The Pollinator Diversity Index was calculated for each population-year datapoint as the Inverse Simpson’s Diversity Index using number of floral visits by pollinator functional group. Inverse Simpson’s Diversity Index is less sensitive to rare occurrences than other indices, and thus was chosen to avoid weighting visits from uncommon visitors (Lazaro et al., 2009). Values of this index were log-transformed to approach normality, though data still include a high number of zeroes, due to datapoints in which only one pollinator functional group was recorded, resulting in a meaningful value of 0 diversity in floral visitors by functional group. We tested for differences in pollinator diversity among species using a GLMM with a Gaussian error distribution, with dataset as a fixed effect and year as a random effect, followed by pairwise multiple comparisons with Tukey adjustments. Due to concerns that these data may be zero-inflated, we examined residuals using package DHARMa for evidence of misspecification, but no significant evidence was identified. Nonetheless, we also performed nonparametric Kruskal-Wallis tests (which are not influenced by potential zero-inflation of data) on population average values of the Pollinator Diversity Index (averaged across dataset type) and present these results as well. Finally, we assessed whether Pollinator Diversity varied geographically by latitude across the ranges of the focal species by performing a GLMM as described above with the addition of a species fixed effect.

### Are certain floral traits associated with visitation from different pollinator groups?

To investigate how floral traits may relate to visitation from different pollinator functional groups across taxonomic boundaries, we performed multiple regression analyses using pooled floral trait data for all four species. We ran a separate GLMM for each pollinator functional group, with proportion of visits as the response variable and population mean values of all seven floral traits as predictors (scaled using the z transformation with the scale() function in tidyverse), along with dataset type as a fixed effect and year as a random effect. Floral trait values were not measured in each year sampled for pollinator visitation, due to time constraints and because measurements were not expected to vary widely among years in these perennial species, so single-year floral trait measurements are taken as population mean values in these models. Models were performed with betabinomial distributions, weighted by total number of visits. Initial models were run with all floral traits, but only terms with p-value > 0.15 were retained in the final model (following Lazaro et al., 2009), though dataset type was retained as a fixed effect.

We also examined whether floral traits impacted the diversity of floral visitors by performing multiple regression analysis with Pollinator Diversity Index (log-transformed) as the response variable. GLMMs were performed as described above except for the use of a Gaussian distribution. As mentioned previously, due to concerns about frequency of zeroes in Pollinator Diversity Index values, we repeated final analyses for significant traits with a single predictor using Kruskal-Wallis tests and report these findings as well.

### Does plant fitness vary in association with visitation from certain pollinator groups?

We tested for variation in fruit set among species using GLMMs with individual-level fruit set as the response and species as the predictor. Models were run with a betabinomial distribution weighted by number of flowers, with population and year included as random effects. Additionally, to assess variation in fruit set across geography, we performed GLMMs as described above using population latitude (a proxy for geography) as a predictor, along with species as a fixed effect and year as a random effect.

Next, we further examined whether population average fruit set varied with respect to visitation from different pollinator groups. For these analyses, data from *C. sessiliflora* and from the *C. purpurea* species complex were treated separately, given observed differences in floral traits and floral visitation between these groups. Individual species of the *C. purpurea* complex were not analyzed separately due to constraints of sample size; however, species was included as a random effect in these models to account for differences among species. We used GLMMs with a betabinomial distribution, weighted by average number of recorded flowers per population, with dataset type as a fixed effect. The response variable was average proportion fruit set for a given population, and predictor was the proportion of visits from a certain pollinator group for data collected in the same year in which fruit set was measured, which resulted in 15 paired-year datapoints for *C. sessiliflora* and 18 for the *C. purpurea* complex. We also tested whether pollinator diversity influenced fruit set of populations, by running GLMMs as described above with Pollinator Diversity Index (Inverse Simpson’s Diversity Index, details above) as the predictor. Finally, we assessed whether populations with greater numbers of overall floral visits experienced increased fruit set by performing GLMMs with total number of floral visits as the predictor term.

## Results

### Floral traits vary among and within species

Our MANOVA revealed significant variation across all floral traits among species (approximate F_3,21,661_= 351.03; p < 0.001) and among populations nested within species (approximate F_19,133,661_= 14.67; p < 0.001). For individual floral traits, all traits varied significantly among species and among populations nested within species (Table S3), and pairwise multiple comparisons revealed variation in key traits between species pairs (Table S4; Supplemental Fig. S2). Overall, the corollas of *Castilleja sessiliflora* were significantly longer than those of the other species, with a prominent floral lip and minimal stigma exertion (Fig. 1; Supplemental Fig. S2). In addition, the bracts subtending these flowers were significantly narrower compared to the other three species and varied in color from green to pale pink from northeast to southwest across the range (Fig. 1B-D), with two distinct populations bearing vivid yellow (Fig. 1E) or pink (Fig. 1F) inflorescences at the southern range extent. Within the *C. purpurea* species complex, *C. lindheimeri* had significantly longer and thinner corollas than either *C. purpurea* and *C. citrina*, along with the most exserted stigmas, and its red bracts were the broadest of all taxa. The purple-bracted *C. purpurea* and yellow-bracted *C. citrina* showed few morphological differences from each other except for corolla length (shortest in *C. citrina* of all species) and corolla width, which was widest in *C. purpurea* (Supplemental Fig. S2; complete statistics Table S3, S4).

Photographs from a subset of populations showed evidence that inflorescences of *C. purpurea* may reflect in the UV spectrum (Supplemental Fig. S3), while *C. lindheimeri, C. citrina*, and the yellow morph of *C. sessiliflora* at SMP did not appear to reflect UV. The pink morph of *C. sessiliflora* at SIC also showed potential reflectance in these UV photos, as did the more typical floral morph of *C. sessiliflora* (photographed at SBL), though the latter was less clear. Previous analysis of typical white-green morphs of *C. sessiliflora* in the northern range found that while the vacuolar pigments of floral tissues did not transmit UV light, hairs on these tissues fluoresced under UV, suggesting the flowers may still reflect at these wavelengths (Crosswhite and Crosswhite, 1970). While preliminary, this evidence suggests that UV reflectance may vary within and between focal species, and additional work should incorporate techniques such as reflectance spectrometry or floral pigment analysis to assess this variation.

### Visitation by pollinator groups varies among and within species

In total, we recorded 4,012 pollinator visits across all sites and years (Table 1). The most common functional groups recorded were hawkmoths (1,161 visits), followed by small and medium bees (1,035 visits). The next most frequent visitors were bumblebees and large bees (670 visits), hummingbirds (637 visits) and finally butterflies (397 visits). Visits from other functional groups (mainly non-Sphingid moths, bee-flies and flies) made up only a small number of visits (112 visits) and were pooled in the “other” category. Analysis of pollinator visitation among species revealed similar patterns for both count and proportion data, and we focus on proportion models hereafter (see Tables S5-S6 for complete results). We found evidence that visitation from hummingbirds and butterflies varied significantly among species (Fig. 3A,B; Table S5), with visitation from small/medium bees showing marginal significance for proportion data (though significant for count models; Table S5). Hummingbirds were the most frequent visitor to *C. lindheimerii* and contributed significantly more visits to this species than to *C. citrina* and *C. purpurea* (Table S6), while zero hummingbirds were observed visiting *C. sessiliflora* (Table 1). Butterflies visited all species at relatively low levels, though *C. purpurea* had significantly higher butterfly visitation than *C. sessiliflora* (Table S6). Small and medium bees were the most common visitor to *C. sessiliflora* and second most common to *C. purpurea* (Table 1; Fig. 3A,B), and proportion of visits from small/medium bees varied marginally among species (Table S5). Although large bees were the most frequent visitor to *C. citrina* and zero large bees were recorded visiting *C. lindheimeri*, differences in their visitation among species was not significant (Table S5). Similarly, hawkmoths were among the most frequent visitors to *C. purpurea* and *C. sessiliflora*, but variation among species was not significant. Lastly, visitation from other visitors did not vary among species for proportion but varied marginally for count data. Dataset effect was significant for models of all pollinator groups except hawkmoths and other visitors (Table S5), which is likely due to the difficulty of seeing small/medium bees at longer ranges and the effect of close observation on foraging behavior of hummingbirds, etc.

**Figure 3.**
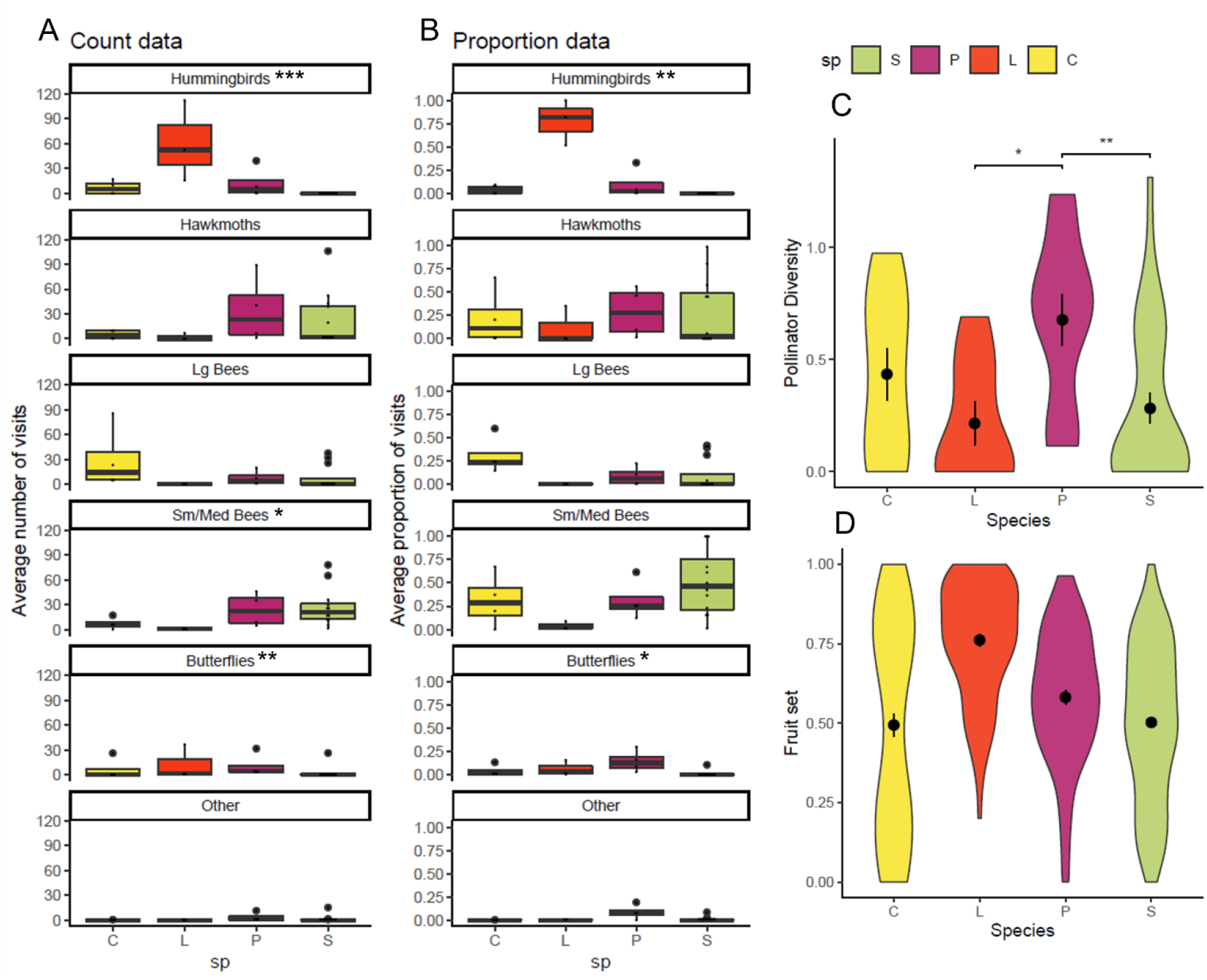
Floral visitation, pollinator diversity, and fruit set by species (sp, C = *C. citrina*, L = *C. lindheimeri*, P = *C. purpurea*, S = *C. sessiliflora*). A) Count data of floral visits (average number of visits) and B) proportions of floral visits (average proportion of visits) from each pollinator functional group by species. Plotted points are population mean values, averaged across years and datasets. Center bars: median value per species; upper and lower hinges: first and third quartiles; whiskers: points within 1.5*IQR of hinges; large points: outlying points (all datapoints are shown as small points overlying plots). C) Pollinator Diversity Index by species. D) Fruit set (fitness) by species (fruit-to-flower ratio for individual plants, N = 591). Violin plots show density of datapoints, with species mean (points) +/- SE (error bars). Asterisks denote significant variation among species (A, B: Tables S5, S6) or pairwise differences among species based on multiple comparisons (C: Table S8) following GLMMs (* 0.05 > p > 0.01; ** 0.01 > p > 0.001; *** 0.001 > p).

For variation within species, we found that all populations of *C. citrina* were visited by bumblebees (Fig. 2), but the contribution of additional visitors varied by site. The most southern population (CHCL) was also visited by butterflies, hawkmoths and hummingbirds, while the northernmost population (CMSC) was visited predominately by hawkmoths and small bees. For *C. lindheimeri*, hummingbirds were the most common visitor to all populations, and the only visitor at one site (LMN), though other populations were also visited by butterflies, hawkmoths, and occasional small bees. Notably, we recorded no visitation by bumblebees/large bees to *C. lindheimeri* during our observation periods. Finally, the species with the most diverse visitors was *C. purpurea*, with at least four of the six pollinator functional groups recorded at every site, and two populations (PMT and PTH) experiencing visitation from all observed pollinator functional groups, which was not observed in any other focal species. Hawkmoths were the most common visitor to *C. purpurea* in the center of the range (PCM and PTH), while small bees were dominant at the northernmost population (PTMS) and hummingbirds at the most southern population (PMT).

Across its wide range, *C. sessiliflora* also exhibited a diversity of visitors (as all functional groups but hummingbirds were recorded), though diversity within populations was low overall (Fig. 2). Hawkmoths were the most frequent visitor at most populations in the southern portion of the range, but no hawkmoths were observed at any northern populations, resulting in a significant decrease in hawkmoth visitation from south to north (Fig. 4; Table S7; χ^2^_1,26_= 11.10, p = 0.001). Small/medium bees were common visitors across the range of *C. sessiliflora* and accounted for most visits in the northern range, which resulted in the proportion of visits from small/medium bees to increase with latitude (χ^2^_1,26_= 14.89, p < 0.001). Two southern populations (SIC and SMP) were previously characterized as having distinctive floral morphs, including shorter corollas and bright inflorescence color (Wenzell et al., 2021), and these populations experienced visitation from a broader array of visitors than other populations of *C. sessiliflora* (Fig. 2). The distinct bright-pink-flowered population (SIC) was the only population of *C. sessiliflora* to experience visitation from butterflies and bee-flies (Bombyliidae; included in “other visitors”), making its assemblage of floral visitors unique compared to other populations of *C. sessiliflora*. Additionally, the yellow-flowered population (SMP) saw visitation by bumblebees in all three years studied, although bumblebee visitation was also observed at two northern white-green-flowered populations in 2019. Latitudinal trends were not significant for any other pollinator functional groups, including bumblebees (χ^2^_1,26_= 1.69 p = 0.19) and other visitors (χ^2^_1,26_= 0.38, p = 0.53), and butterflies (recorded at only one population) and hummingbirds (not recorded at any *C. sessiliflora* sites) were too infrequent for analysis (Fig. 4; Table S7). Dataset effect was significant for small/medium bees (χ^2^_1,26_= 12.4, p < 0.001) and large bees/bumblebees (χ^2^_1,26_= 7.43 p = 0.006), but not for any other functional groups, again likely due to the difficulty observing small bees at longer visual ranges.

**Figure 4.**
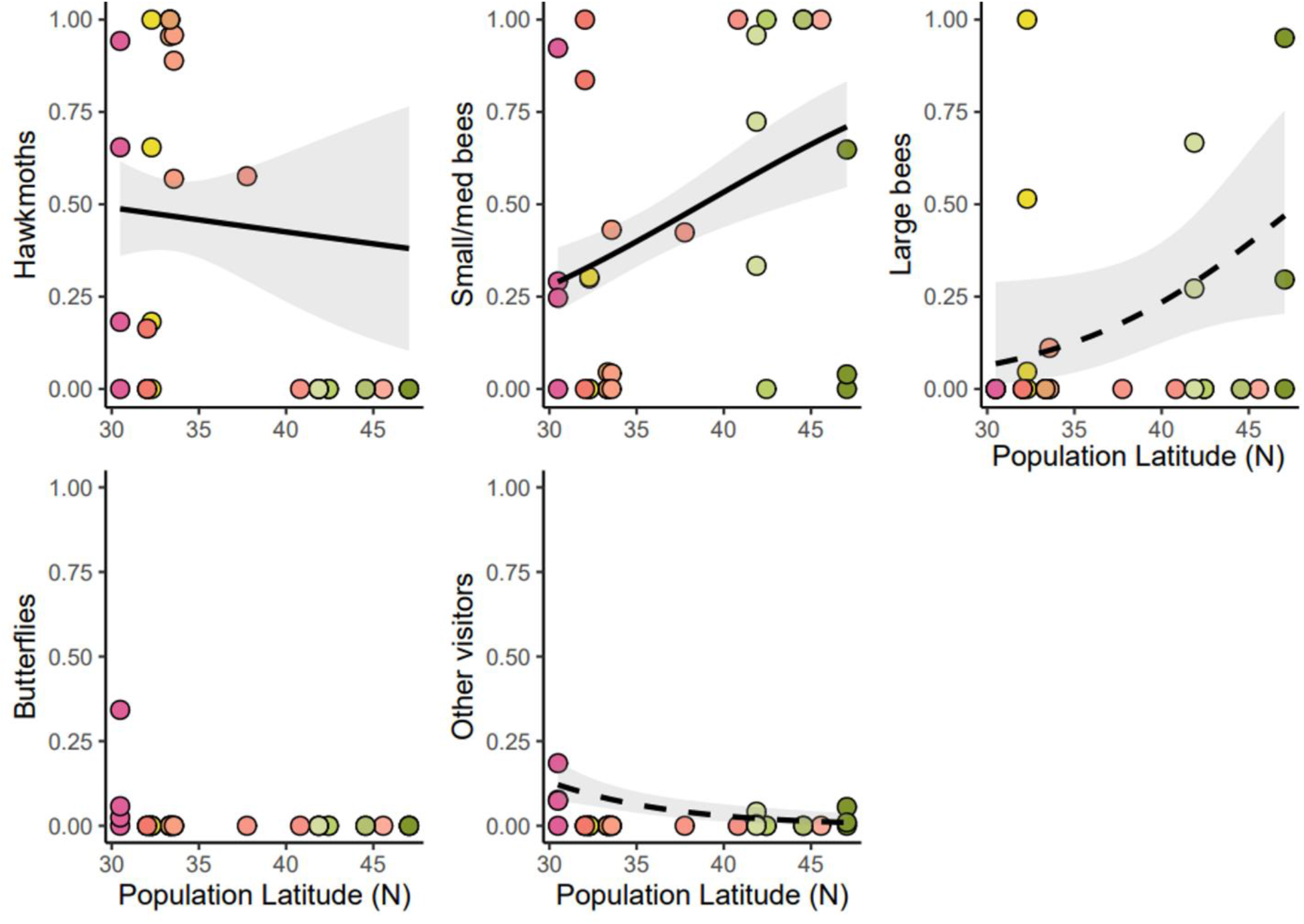
Geographic variation in visitation across the range of *C. sessiliflora*. Proportion visitation by pollinator functional group (population-year-dataset datapoints, N = 31) is plotted against population latitude (° N). Fill colors show median floral color per population. Solid trend lines denote a significant relationship (p < 0.05); dashed lines denote nonsignificant (p > 0.05) based on GLMM; no trend line indicates analyses were not supported. Note no hummingbirds were observed visiting *C. sessiliflora*.

Diversity of floral visitors (measured as Pollinator Diversity Index) varied among species (Fig. 3C) according to both the GLMM (species: χ^2^_3,50_= 12.48, p = 0.006; dataset: χ^2^_1,50_= 0.12, p = 0.73) and the Kruskal-Wallis rank sum test of population average values (χ^2^_3,22_ = 8.8, p = 0.03). Multiple comparisons of the GLMM revealed that *C. purpurea* exhibited a significantly higher diversity of floral visitors compared to *C. lindheimeri* (t_50_ = -2.92, p = 0.026) and *C. sessiliflora* (t_50_ = 3.18, p = 0.013). Finally, we did not find evidence that pollinator diversity varied significantly by latitude (Supplemental Fig. S4A) based on the GLMM (latitude: χ^2^_1,49_ = 1.73, p = 0.19; species: χ^2^_3,49_= 12.14, p = 0.007; dataset: χ^2^_1,49_= 0.11, p = 0.73).

### Visitation from different pollinator groups is associated with floral traits

We identified relationships between floral traits and visitation from different pollinator groups across species (Table 2). Visitation from hummingbirds was positively associated with greater population mean corolla length and stigma exsertion, but negatively associated with length of petaloid lips. For hawkmoths, visitation was associated with wider corollas, shorter lips, and narrower bract lobes. Bumblebee/large bee visitation was negatively associated with NMDS2 axis of floral color, consistent with greater visitation to human-vision yellow-green flowers than to red/purple flowers (Fig. 5A). Visitation from small/medium bees was positively associated with longer lips and wider bract lobes but negatively associated with stigma exsertion and corolla length. For butterflies, visitation was associated with shorter corollas and higher values on the NMDS2 color axis, in line with red-purple floral color (Fig. 5B). Visitation from other visitors was associated with narrower corollas, wider bract lobes, and higher values on the NMDS1 color axis (associated with human-vision purple and pink), though this group of “other visitors” includes various functional and taxonomic groups (e.g., bee flies, flies, and moths) expected to respond to floral signals in differing ways. Dataset type was significant for all pollinator groups excepts hawkmoths and other visitors.

**Table 2.**
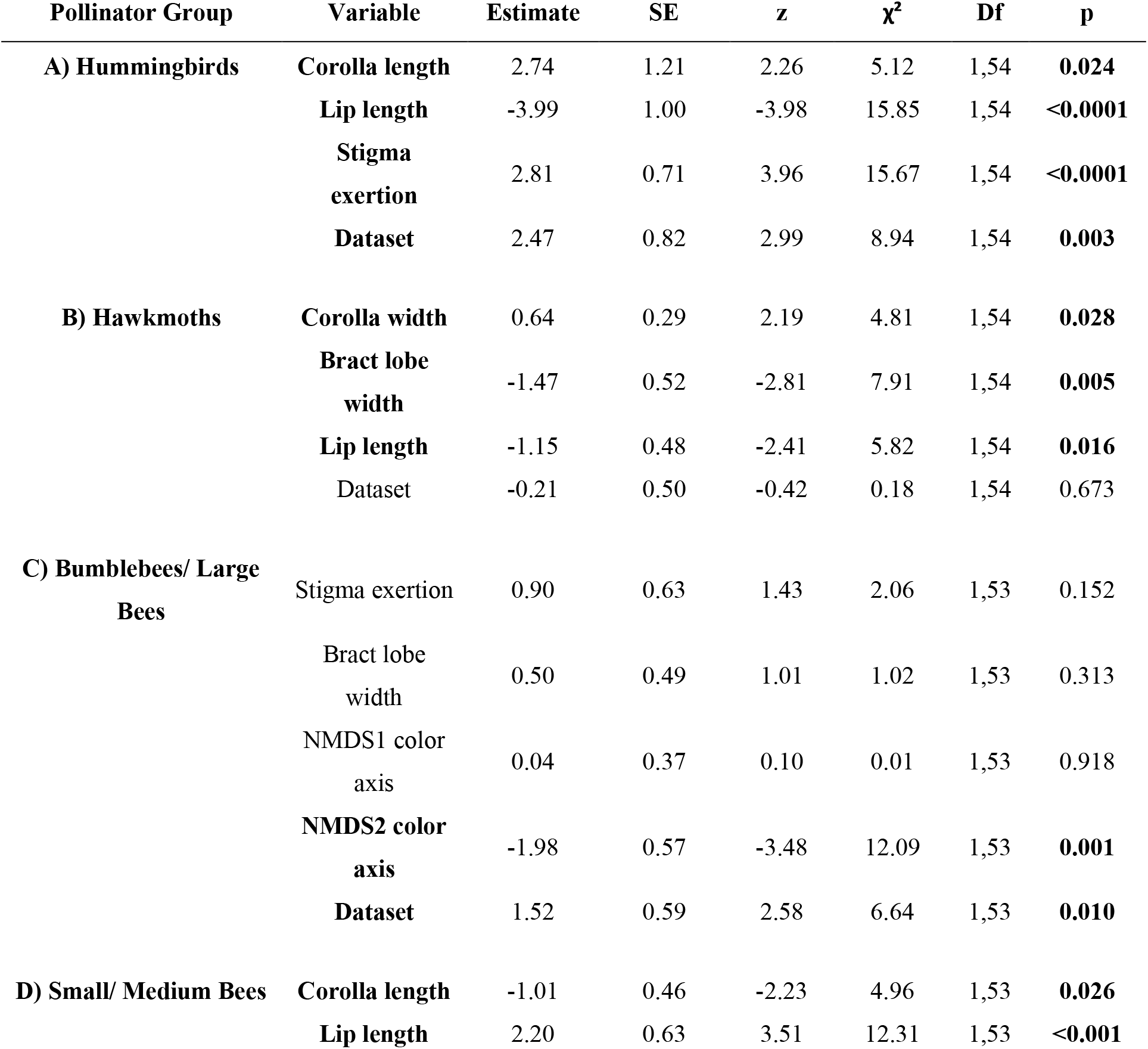
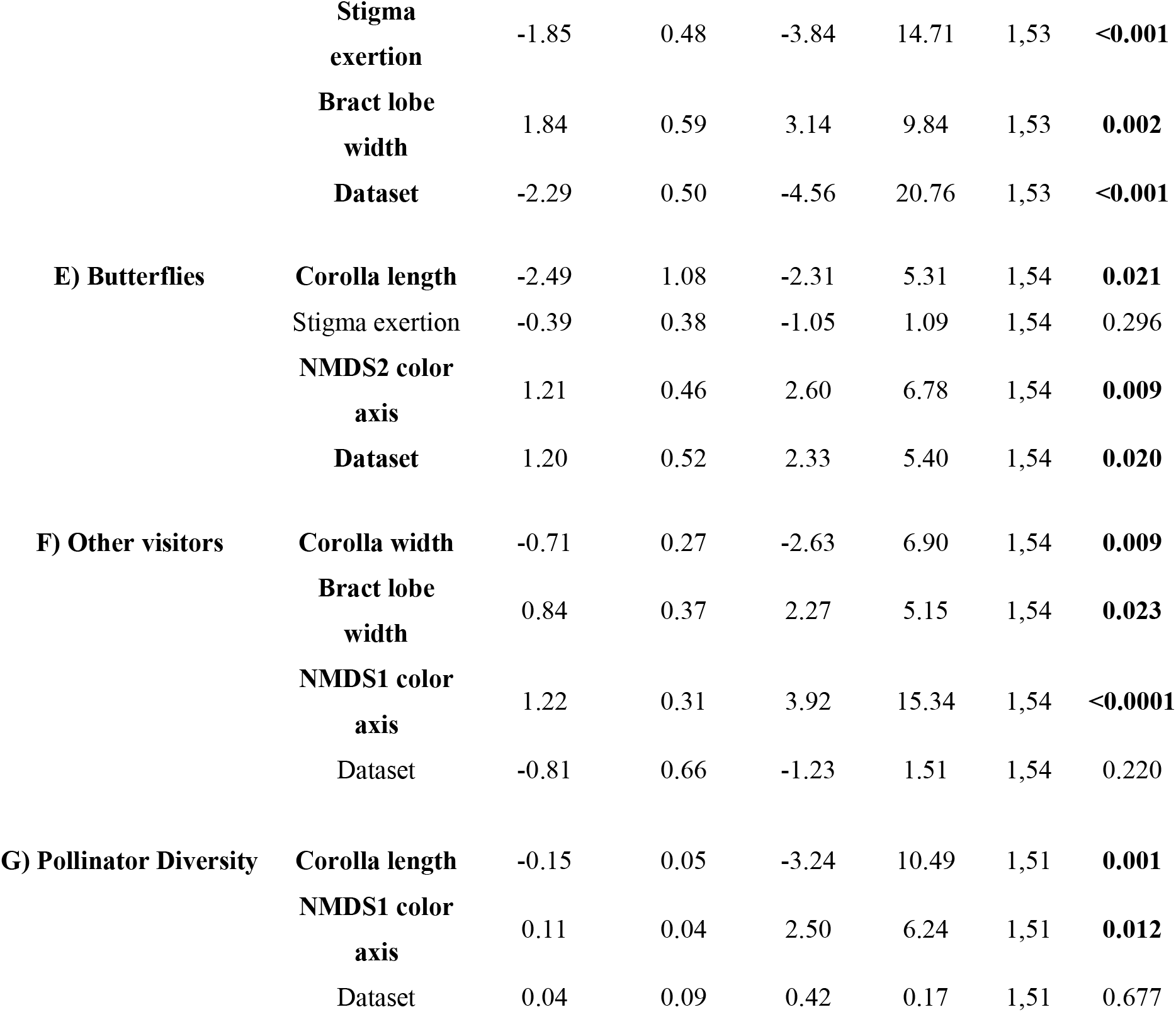
Visitation by pollinator group in relation to floral trait values. Final GLMMs of scaled population-average floral trait values with visitation from different pollinator groups (A-F) and Pollinator Diversity (G). Models were run for each pollinator group (proportion of visits)/ diversity metric and initially included all 7 floral traits (see text for details). Final models included only variables with a p-value > 0.15 in the initial model. Dataset type was included as a fixed effect in all models, and year was included as a random effect. Coefficient estimate (Estimate, in logit scale for A-F), standard error (SE), z-value (z) and p-value (p) are presented for each variable, along with **χ**^**2**^ and degrees of freedom (Df, with residual Df). P < 0.05 and associated variables are in bold.

| Pollinator Group | Variable | Estimate | SE | z | $\chi^2$ | Df | p |
| --- | --- | --- | --- | --- | --- | --- | --- |
| **A) Hummingbirds** | **Corolla length** | 2.74 | 1.21 | 2.26 | 5.12 | 1,54 | **0.024** |
|  | **Lip length** | -3.99 | 1.00 | -3.98 | 15.85 | 1,54 | **<0.0001** |
|  | **Stigma exertion** | 2.81 | 0.71 | 3.96 | 15.67 | 1,54 | **<0.0001** |
|  | **Dataset** | 2.47 | 0.82 | 2.99 | 8.94 | 1,54 | **0.003** |
| **B) Hawkmoths** | **Corolla width** | 0.64 | 0.29 | 2.19 | 4.81 | 1,54 | **0.028** |
|  | **Bract lobe width** | -1.47 | 0.52 | -2.81 | 7.91 | 1,54 | **0.005** |
|  | **Lip length** | -1.15 | 0.48 | -2.41 | 5.82 | 1,54 | **0.016** |
|  | Dataset | -0.21 | 0.50 | -0.42 | 0.18 | 1,54 | 0.673 |
| **C) Bumblebees/ Large Bees** | Stigma exertion | 0.90 | 0.63 | 1.43 | 2.06 | 1,53 | 0.152 |
|  | Bract lobe width | 0.50 | 0.49 | 1.01 | 1.02 | 1,53 | 0.313 |
|  | NMDS1 color axis | 0.04 | 0.37 | 0.10 | 0.01 | 1,53 | 0.918 |
|  | **NMDS2 color axis** | -1.98 | 0.57 | -3.48 | 12.09 | 1,53 | **0.001** |
|  | **Dataset** | 1.52 | 0.59 | 2.58 | 6.64 | 1,53 | **0.010** |
| **D) Small/ Medium Bees** | **Corolla length** | -1.01 | 0.46 | -2.23 | 4.96 | 1,53 | **0.026** |
|  | **Lip length** | 2.20 | 0.63 | 3.51 | 12.31 | 1,53 | **<0.001** |
|  | **Stigma exertion** | -1.85 | 0.48 | -3.84 | 14.71 | 1,53 | **<0.001** |
|  | **Bract lobe width** | 1.84 | 0.59 | 3.14 | 9.84 | 1,53 | **0.002** |
|  | **Dataset** | -2.29 | 0.50 | -4.56 | 20.76 | 1,53 | **<0.001** |
| **E) Butterflies** | **Corolla length** | -2.49 | 1.08 | -2.31 | 5.31 | 1,54 | **0.021** |
|  | Stigma exertion | -0.39 | 0.38 | -1.05 | 1.09 | 1,54 | 0.296 |
|  | **NMDS2 color axis** | 1.21 | 0.46 | 2.60 | 6.78 | 1,54 | **0.009** |
|  | **Dataset** | 1.20 | 0.52 | 2.33 | 5.40 | 1,54 | **0.020** |
| **F) Other visitors** | **Corolla width** | -0.71 | 0.27 | -2.63 | 6.90 | 1,54 | **0.009** |
|  | **Bract lobe width** | 0.84 | 0.37 | 2.27 | 5.15 | 1,54 | **0.023** |
|  | **NMDS1 color axis** | 1.22 | 0.31 | 3.92 | 15.34 | 1,54 | **<0.0001** |
|  | Dataset | -0.81 | 0.66 | -1.23 | 1.51 | 1,54 | 0.220 |
| **G) Pollinator Diversity** | **Corolla length** | -0.15 | 0.05 | -3.24 | 10.49 | 1,51 | **0.001** |
|  | **NMDS1 color axis** | 0.11 | 0.04 | 2.50 | 6.24 | 1,51 | **0.012** |
|  | Dataset | 0.04 | 0.09 | 0.42 | 0.17 | 1,51 | 0.677 |

**Figure 5.**
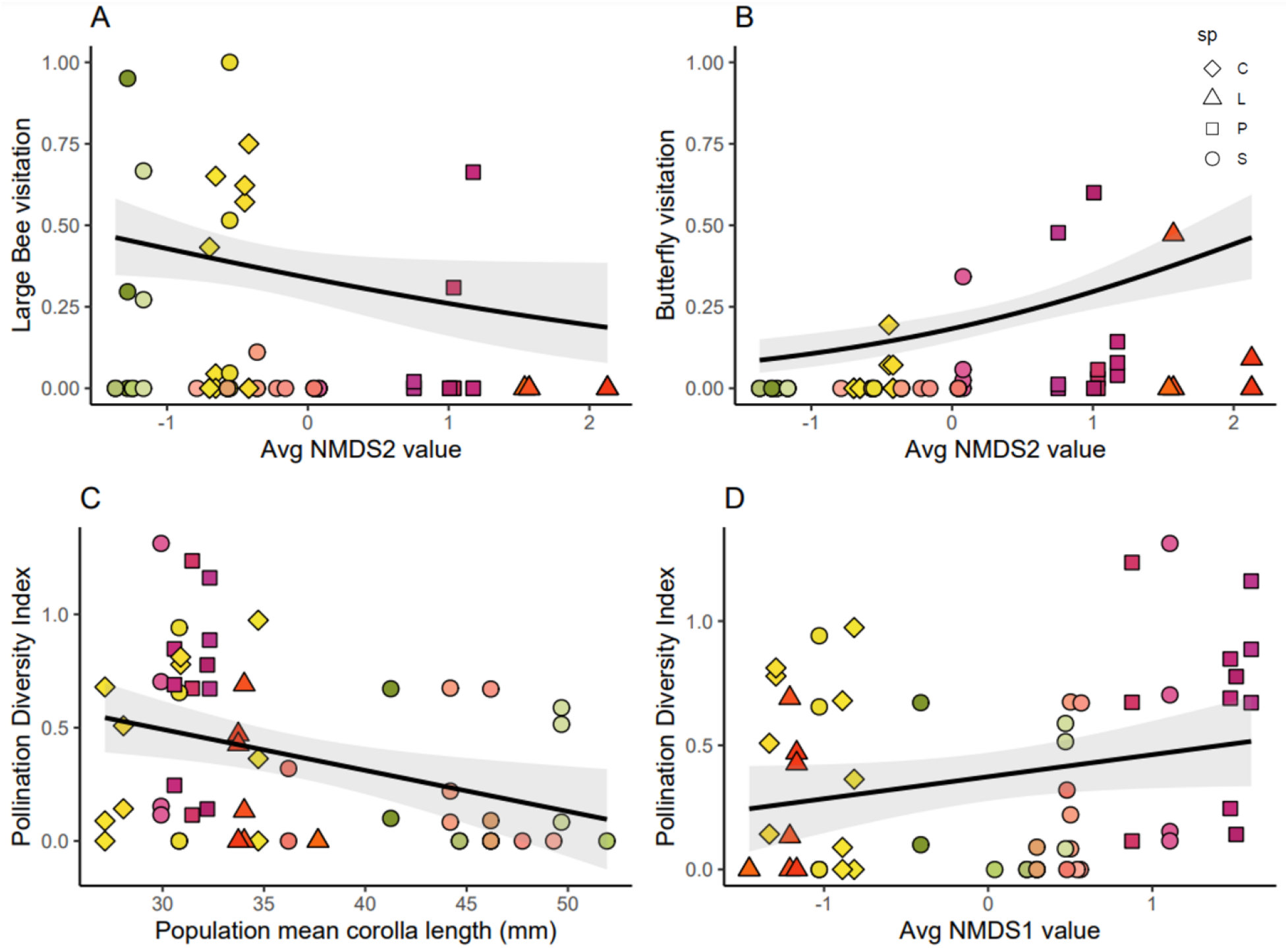
Pollinator visitation and diversity by floral trait values. Top row: population mean NMDS2 axis value (floral color) in relation to proportion of visits from A) large bees/bumblebees and B) butterflies. Bottom row: floral trait values C) population mean corolla length and D) population mean NMDS1 axis value (floral color) in relation to pollinator diversity index. Points show species (sp): *C. citrina* (diamonds), *C. lindheimeri* (triangles), *C. purpurea* (squares), *C. sessiliflora* (circles), and fill color shows population median floral color. Trend line represents significant relationships (p < 0.05) based on GLMM (Table 2).

Pollinator diversity was significantly associated with two traits (Table 2G): corolla length was negatively associated, indicating that populations with longer corollas were associated with less diverse visitors (Fig. 5C), while average NMDS1 color value was positively associated, corresponding to human-vision cool pink and purple flowers (Fig. 5D). Because the response variable data had a large number of zeroes, we also ran a Kruskal-Wallis rank sum test of pollinator diversity against population mean corolla length and mean NMDS1 value, both of which were not significant (corolla length: χ^2^_22,56_ = 29.33, p = 0.14; NMDS1: χ^2^_22,56_ = 27.51, p = 0.12).

### Plant fitness increases with visitation from certain pollinator groups

Fruit set did not vary significantly among species (Fig. 3D; χ^2^_3,584_ = 6.25, p = 0.099) when a population random effect was included (suggesting that variation at the population level may outweigh that among species), but fruit set did vary geographically, decreasing at greater latitudes (Supplemental Fig. S4B; χ^2^_1,584_ = 12.63, p = 0.0004) with a significant species term (χ^2^_1,584_ = 65.14, p < 0.0001).

Next, we found evidence that visitation from certain pollinator groups was associated with population-level fruit set within *C. sessiliflora* and within the *C. purpurea* complex (Fig. 6). Populations of *C. sessiliflora* with a higher proportion of visits from hawkmoths had higher average fruit set (Fig. 6A; Table S9). In contrast, neither the proportion of visits from small/medium bees nor bumblebees/large bees (the most common other visitors to *C. sessiliflora*) were a significant predictor of fruit set (Fig. 6C; Table S9). Butterflies and other visitors were uncommon visitors to *C. sessiliflora*, and neither of these groups was significantly associated with population average fruit set (Table S9). Within the *C. purpurea* complex, increased proportion of visits from hummingbirds was associated with higher average fruit set (Fig. 6B; Table S9), while no significant relationship was found between average fruit set and proportion of visits from small/medium bees (Fig. 6C) or from hawkmoths (Fig. 6A; Table S9). Relationships between fruit set and visitation from bumblebees/large bees, butterflies, and other visitors were not significant (Table S9) and were not common visitors in the paired-year datapoints used for these analyses.

**Figure 6.**
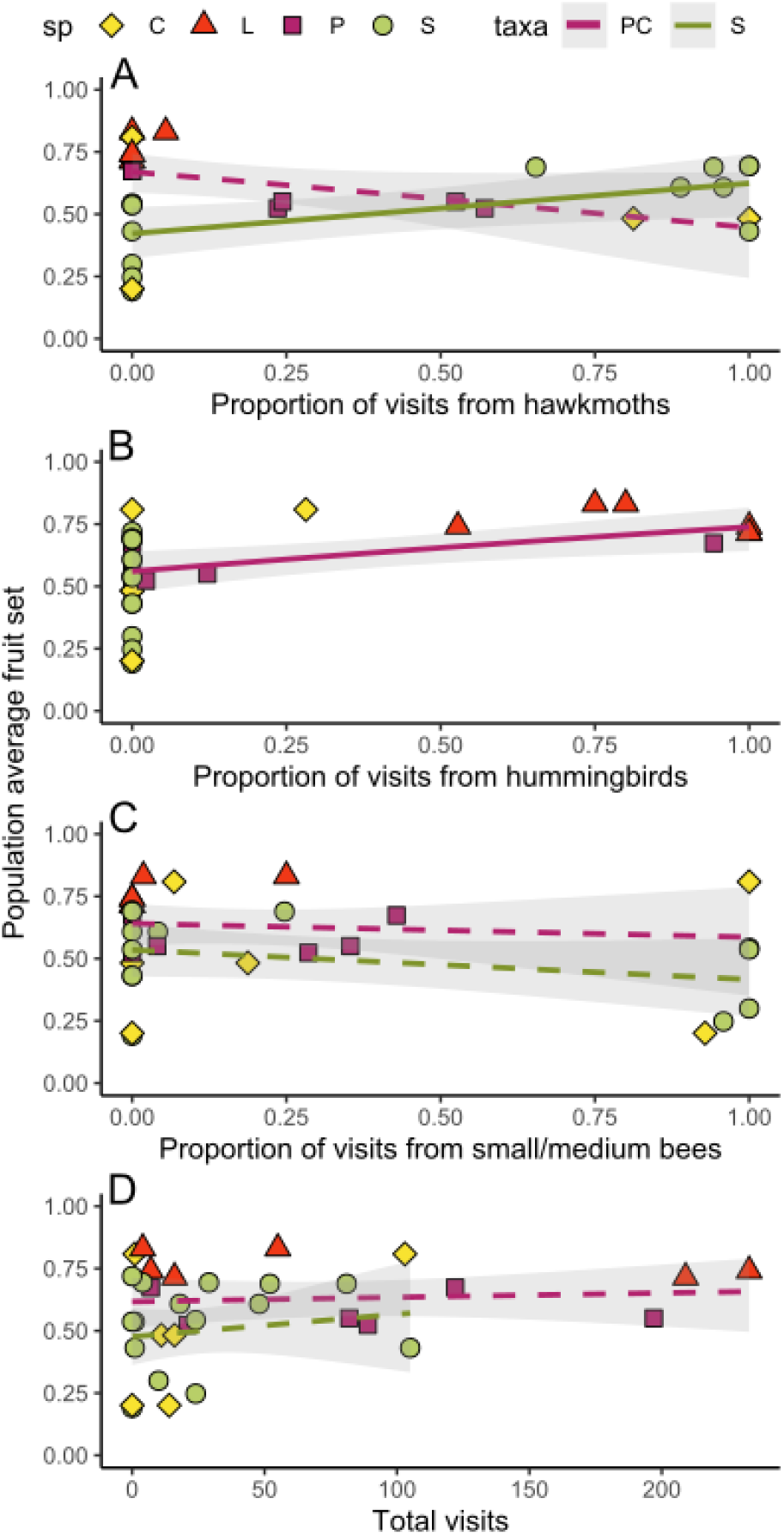
Population fitness by pollinator visitation. Population average fruit set is plotted against proportion of visits from certain pollinator groups (A: hawkmoths, B: hummingbirds, C: small/medium bees) and D) total number of visits to that population during the same calendar year for *C. sessiliflora* (green trend line) and the species of the *C. purpurea* complex (purple trend line). Points are coded by species: C = *C. citrina*, L = *C. lindheimeri*, P = *C. purpurea*, S = *C. sessiliflora*. Solid trend lines denote a significant relationship (p < 0.05); dashed lines denote nonsignificant relationship (p > 0.05) based on GLMM (Table S9).

Furthermore, we did not find evidence for a significant relationship between pollinator diversity value and fruit set in either *C. sessiliflora* or the *C. purpurea* complex (Table S9). Finally, to assess whether these findings were attributable to specific pollinator groups per se, or merely reflected an increase in total visitation yielding increased fruit set, we also tested whether total number of floral visits influenced fruit set. These models did not find evidence that total number of floral visits (regardless of pollinator identity) was associated with fruit set in either *C. sessiliflora* or the *C. purpurea* complex (Fig. 6D; Table S9). The dataset fixed effect was not significant in any models (Table S9).

## Discussion

In this study we characterize geographic variation in pollinator visitation and reproductive fitness across the range of four species of *Castilleja*. By sampling multiple populations over several years, we document a wide spectrum of visitors spanning hummingbirds, Lepidoptera (butterflies and hawkmoths), Hymenoptera (mostly small solitary bees and bumblebees) and Diptera (bee-flies and occasional hoverflies), demonstrating a diversity of visitors consistent with studies of other *Castilleja* species (Hersch and Roy, 2007; Hilpman and Busch, 2021). Despite exhibiting significant divergence in floral traits, we show that our four focal *Castilleja* species were visited broadly by all pollinator groups (with two exceptions: the lack of hummingbird visits to *C. sessiliflora* and of large bee visits to *C. lindheimeri*). Despite this diversity, we found that visitor assemblages at the population level represented a mosaic of interactions likely to contribute to observed floral trait variation. Furthermore, we present evidence that visitation from different pollinator functional groups is associated with variation in floral traits, suggesting local pollinators may play a role in shaping floral variation. We also found that greater visitation from certain pollinator groups correlated with increased reproductive fitness, indicating pollinators may be capable of exerting selection. Hence, despite the wide diversity of visitors observed across species, consistent with a generalized pollination mode (e.g., Waser et al., 1996; Ohashi et al., 2021), we find multiple lines of evidence to support the hypothesis that a geographic mosaic of pollinators could be mediating divergence in floral traits in these recently diverged plant species.

### *Pollinator mosaics among the species of the* Castilleja purpurea *complex*

We explored whether variation in floral visitors underlies floral divergence among species within the *C. purpurea* species complex. These species exhibited divergence in floral color across narrow geographic clines despite low genetic differentiation (Wenzell et al., 2021), which is consistent with other examples of pollinator-mediated selection driving floral color transitions (Streisfeld and Kohn, 2005; Hopkins et al., 2012; Stankowski et al., 2017). By investigating whether adaptation to distinct pollinators could be a driver of floral divergence, we found compelling support for this hypothesis in the predominately hummingbird-pollinated *C. lindheimeri*, some support in the largely bee-pollinated *C. citrina*, but less clear evidence in *C. purpurea*, which hosted a wide diversity of visitor groups, consistent with a generalist pollination mode.

Although the species of the *C. purpurea* complex were visited by a broad array of pollinator functional groups overall, patterns varied by species. Within the red-bracted *C. lindheimeri*, which had low pollinator diversity and was visited predominately by hummingbirds at all sampled populations, several lines of evidence suggest possible floral adaptation to hummingbird pollinators. First, hummingbirds visited *C. lindheimeri* significantly more frequently than any other species, and visitation by hummingbirds was associated with longer corollas, more exserted stigmas, and shorter petaloid lips, all of which are floral traits associated with *C. lindheimeri* flowers (Supplemental Fig. S2) and with hummingbird pollination modes (Faegri and van der Pijl, 1971; Fenster et al., 2004; Rosas-Guerrero et al., 2014). *Castilleja lindheimeri* also exhibits red floral pigmentation, another well-documented hummingbird-associated trait, though our model found no association between hummingbird visitation and color values. While surprising, this finding may reflect the hypothesis that red pigment in hummingbird-pollinated flowers may function as a deterrent to bee pollinators if not as an attractant to hummingbirds, per se (e.g., Castellanos et al., 2004, Schiestl and Johnson, 2013; discussed further below). Finally, higher visitation by hummingbirds was associated with greater average fitness (fruit set) of populations (Fig. 6B), which suggests hummingbird pollinators may exert selection on flowers of *C. lindheimeri* through increased reproductive fitness. However, because these results reflect maternal reproductive fitness (fruit set), which has been shown to vary with the resource richness of a plant’s environment (Harder and Routley, 2007), we cannot rule out that resource availability or other abiotic or biotic factors may also contribute to these patterns. Nonetheless, this result was replicated across multiple sites and was specific to hummingbird visitation (compared to other pollinator groups and total number of visits), thus lending support to our interpretation. While additional work is needed to directly test for pollinator-mediated selection on floral traits in this system, we present evidence consistent with the hypothesis that *C. lindheimeri* may exhibit adaptation to pollination by hummingbirds, based on its suite of syndrome-aligned floral traits and evidence linking hummingbird visitation with increased fitness.

In contrast, visitor assemblages of the other species of the *C. purpurea* complex, *C. purpurea* and *C. citrina*, were more generalized. In terms of floral divergence, this species pair appears to differ less in morphological traits compared to the other focal species (Supplemental Fig. S2). Lower floral divergence compared to other focal species may be expected with generalist pollinator assemblages, which may exert contrasting selection pressures on floral traits, resulting in intermediate phenotypes (van der Niet et al., 2014; Ohashi et al., 2021). Nonetheless, some trends in visitation differ between these species. Notably, the yellow-flowered *C. citrina* showed moderate pollinator diversity but was visited most frequently by bumblebees/large bees and was the only species visited by large bees at every population (Figs. 2, 3). We hypothesize that color may play a role in this pattern, given that bumblebee visitation was associated with lower values on the NMDS2 color axis, which includes mostly human-vision yellow and green inflorescences (Fig. 5A). Additionally, the yellow-flowered population of *C. sessiliflora* (SMP) had greater visitation by bumblebees compared to other nearby *C. sessiliflora* populations with differing color (Fig. 2). The association between large bee visitation and NMDS2 color value suggests both that large bees are more likely to visit yellow flowers and/or less likely to visit red flowers (Fig. 5A). Thus, this finding may also be consistent with the evolution of red floral pigmentation as a deterrent to bee visitors (Castellanos et al., 2004) given the lower sensitivity of bee visual systems to human-vision red spectra (Schiestl and Johnson, 2013), which may or may not occur in concert with increased attraction of bees to yellow flowers.

*Castilleja purpurea* had the greatest diversity of floral visitors of any species (Fig. 3C), and its most common visitor varied from small/medium bees to hawkmoths to hummingbirds across the range (Fig. 2). Despite this diversity, this species experienced frequent visitation from Lepidopteran pollinators, with significantly greater butterfly visitation than other species, and a high number of visits by hawkmoths. We found that butterfly visitation increased along the NMDS2 axis of color variation linked to human-vision red-purple flower color (Fig. 5B), which is consistent with innate preference for and spectral sensitivity to human-vision red and purple flowers reported in Lepidoptera (Lunau and Maier, 1995; Chittka and Thomson, 2001). While a possible relationship between butterflies and purple and bumblebees and yellow flowers in this system remains largely speculative, these findings warrant future work into floral pigmentation and its evolutionary origins in this species complex.

Our findings suggest flower color may not constitute a strong barrier to pollination by different visitor groups but could nonetheless impact their frequency of visitation. This observation supports suggestions that color is a weak barrier compared to other syndrome-associated floral traits (Dellinger, 2020), though color nonetheless can evolve rapidly and facilitate reproductive isolation through pollinator preference (Schemske and Bradshaw, 1999; Bradshaw and Schemske, 2003) and reinforcement via pollinator foraging behavior (Hopkins and Rausher, 2012). As no clear pollinator shift was observed to explain the color transition between *C. purpurea* and *C. citrina*, future work should investigate floral antagonists as potential selective agents on color in this system, given their important role in exerting selection on floral traits (Irwin et al., 2003; Kessler et al., 2010; Jogesh et al., 2017). Taken together, this study provides support for the hypothesis that variable pollinator assemblages likely contribute to floral trait divergence among species of the *C. purpurea* species complex, though additional work is needed to characterize other potential ecological drivers.

### *Pollinator mosaics across the range of* Castilleja sessiliflora

Compared to its congeners, flowers of *C. sessiliflora* are clearly differentiated by long corolla tubes, long petaloid lips, and pale floral pigmentation (Fig. 1). Beyond this, *C. sessiliflora* exhibits intraspecific floral trait variation, including latitudinal variation in inflorescence color across its range and more divergent floral morphs in the south, which vary in color and corolla length (Fig. 1; Wenzell et al., 2021) and which occur in regions adjacent to members of the *C. purpurea* complex species (discussed below). In our study, floral trait variation across the distribution of *C. sessiliflora* corresponded to range-wide variation in floral visitors. Visitation to northern populations came almost exclusively from small/medium bees and less commonly bumblebees, which were previously reported to visit *C. sessiliflora* in Wisconsin (Crosswhite and Crosswhite, 1970). In the southern portion of the range, the most common visitors were hawkmoths (*Hyles lineata*) and small/medium bees, along with occasional bumblebees, butterflies, and other infrequent visitors (e.g., bee-flies) at a few populations (Fig. 2). Given that hawkmoths were the predominant visitor to many populations in the south, it was noteworthy that no hawkmoths were observed foraging on *C. sessiliflora* in the northern half of the range, despite reports of them at some study sites (Friends of Nachusa Grasslands, 2020) and rare visits to *C. sessiliflora* observed in previous years (J. Fant, unpublished data). Hawkmoth visitation is known to be unreliable in space and time (Miller, 1981; Campbell et al., 1997), and it is possible that hawkmoths in the northern region may be active outside our observation window (e.g., later at night) or later in the growing season than the spring-flowering *C. sessiliflora*. Additionally, because species were sampled unevenly across years (Table 1; Table S1), we cannot rule out that potential temporal fluctuations in visitor assemblages may influence observed differences among species. However, reported patterns were consistent across multiple populations and regions sampled in multiple years, which lends support to out interpretations.

Within *C. sessiliflora*, we identified two phenotypically distinct populations that were visited by broad pollinator assemblages that may more closely resemble those of the *C. purpurea* complex than other populations of *C. sessiliflora:* the yellow-flowered SMP (Fig. 2D) and pink-flowered SIC (Fig. 2E-F), whose visitors included bumblebees, butterflies, and other visitors in addition to hawkmoths and small bees. Previous genetic analyses did not find evidence that these distinct populations represent recent hybrids with the *C. purpurea* species (Wenzell et al., 2021), suggesting that phenotypic commonalities between these groups could reflect responses to similar ecological pressures. In fact, these two distinct populations of *C. sessiliflora* and most sampled populations of the *C. purpurea* complex occur in dry grasslands throughout Texas in the south-central United States, where a greater apparent abundance and diversity of pollinator functional groups (e.g., butterflies, bumblebees, and hummingbirds) were observed compared to populations of *C. sessiliflora* elsewhere in the range. Thus, these patterns of co-occurring diversity of floral phenotype and of floral visitors are consistent with a geographic mosaic of plant-pollinator interactions, whereby diversity in local pollinators may beget diversity in floral divergence. We hypothesize that these distinct floral phenotypes may attract (via brighter floral colors) and allow access to (via shorter corollas) a more diverse assemblage of visitors than do more typical *C. sessiliflora* flowers (e.g., Fig. 1B-D), which is supported by our finding that vivid color and shorter corollas are associated with increased pollinator diversity (Fig. 5C, D). In such a case, these distinct floral morphs may represent pollination ecotypes within *C. sessiliflora*, which could suggest early stages of speciation, potentially mediated by geographic mosaics of pollinators (van der Niet et al., 2014). This hypothesis should be investigated further in this system by comparing pollination effectiveness of different functional groups, in addition to visitation.

In the northern portion of the range, visitation was often low (Table S2), and fruit set decreased with increasing latitude. This could suggest that small/medium bees, the predominate visitors in the north, may be relatively ineffective pollinators of *C. sessiliflora*. We found that a higher proportion of visits from small/medium bees did not translate to increased fruit set, possibly because small/medium bees, which were only observed foraging on pollen (and did not enter corollas to access nectar), may be small enough to do so without contacting stigmas (see Fig. 2A), potentially resulting in low pollen deposition and poor pollination. Interestingly, visitation from small/medium bees was associated with lower stigma exsertion (Table 2), which could be selected for to facilitate pollination by small bees foraging on anthers. Visitation by small/medium bees was also associated with longer petaloid lips and wider bract lobes, both of which could serve as landing platforms for bees foraging for pollen at the mouths of corollas. Despite the prevalence of small/medium bees and the absence of hawkmoths among northern populations, these populations still exhibit clear hawkmoth-associated floral traits, namely long corolla tubes and pale floral pigmentation (Faegri and van der Pijl, 1971; Fenster et al., 2004). Though further work is needed, we hypothesize this may reflect factors such as evolutionary constraints (e.g., Zufall and Rausher, 2004; Huang and Fenster, 2007), effects of land use change on pollinator populations (Grixti et al., 2009; Young et al., 2017; Durant and Otto, 2019), and/or the history of glaciation in the northern Great Plains and recent northward colonization of plant populations, which may have become adapted to hawkmoth population in their southern range extent (Clayton and Moran, 1982; Waters et al., 2013; Ursenbacher et al., 2015).

Despite the broad assemblage of visitors observed range-wide, we hypothesize hawkmoths may be the most efficient and effective pollinators of *C. sessiliflora* when they are present. Both hawkmoth visitation and fruit set decreased to the north, and populations with greater visitation from hawkmoths experienced higher average fruit set, suggesting hawkmoths may confer higher fitness to those populations, which could drive floral adaptation. Hawkmoths (*Hyles lineata*) have been reported as infrequent but impactful pollinators in *Ipomopsis*, where they exert strong selection on flowers in years they visit (Campbell et al., 1997; Campbell and Aldridge, 2006), thus influencing floral trait evolution even as inconsistent pollinators. In contrast to *C. sessiliflora*, we did not find evidence that increased hawkmoth visitation was associated with increased fruit set in the *C. purpurea* complex, though hawkmoths still visited these species at similar levels. Similarly, despite expected associations between long corolla tubes and pollination by hawkmoths, hawkmoth visitation was not associated with corolla length (Table 2). While unexpected, we suspect this reflects the ability of hawkmoths to forage on flowers of all focal species, as their long proboscides can access nectar in both short and long corollas (Johnson et al., 2017). However, this finding does not contradict the hypothesis that longer corollas may improve the efficiency of pollination by hawkmoths, by increasing contact between plant sexual organs and hawkmoths’ bodies (Whittall and Hodges, 2007). In fact, a pollinator exclusion experiment found evidence that fruit set was greater among plants exposed to nocturnal pollination, but only in long-tubed populations where hawkmoths were present, suggesting both pollinator identity and floral traits mediate increased plant fitness (Wenzell et al., in prep.). Combined with the results presented here, these findings lead us to hypothesize that the long corollas of *C. sessiliflora* may represent an adaptation to pollination by hawkmoths, which warrants further study.

## Conclusions

This study characterizes variation in pollinator assemblages at multiple populations across the ranges of *C. sessiliflora* and the species of the *C. purpurea* complex. Aided by our thorough sampling of 23 populations across the ranges of our focal species, we found considerable variation in floral visitors among species and throughout their geographic ranges, coinciding with variation in floral traits and plant fitness. Within the *C. purpurea* species complex, *C. purpurea* and *C. citrina* were visited by a diverse array of generalist pollinator functional groups. In contrast, *C. lindheimeri* was predominately visited by hummingbirds, was characterized by floral traits associated with hummingbird pollination, and demonstrated increased fruit set at populations with a higher proportion of visits from hummingbirds, suggesting that *C. lindheimeri* may exhibit adaptation to hummingbird pollinators. In *C. sessiliflora*, floral visitors were structured by geography, with hawkmoths being frequent visitors in the southern half of the range, though absent in the north. Despite their inconsistency across the range, hawkmoth visitation was associated with increased fitness to populations of *C. sessiliflora*, which suggests they may be capable of driving adaptation in floral traits through increased plant fitness. Taken together, these results provide evidence that mosaics of pollinators likely play a role in floral divergence in *C. sessiliflora* and the species of the *C. purpurea* complex, though likely in concert with other ecological drivers. Thus, this study provides a robust investigation of how geographic variation in pollinator mosaics across the distributions of plant species may contribute to divergence in floral traits in a recently radiating genus, potentially via pollinator-mediated plant evolution.

## Acknowledgements

The authors thank A. Iler, H. Briggs, K. Byers, M. Neequaye, and two anonymous reviewers for feedback on previous versions of this manuscript. Additional thanks to A. Iler for input on statistical analyses and to K. Byers for assistance with identification of photographed insects. We also thank D. Tank and M. Egger for insight on the *C. purpurea* species complex. We are grateful for field assistance from S. Deans, K. Bonefont, A. Cisternas Fuentes, C. Woolridge, A. Gruver, E. Lewis, S. Todd, A. J. Morgan, K. Manion, M. Malone, and G. Perez Cartagena, and laboratory assistance from J. Zhang. We thank the following permitting agencies for permission and access to study sites: The Nature Conservancy Chapters of TX, OK, and IL, Native Prairies Association of Texas, Cities of Lubbock, TX, Ft. Worth, TX, and Tulsa, OK, TX Parks and Wildlife Department, NM Energy, Minerals and Natural Resources Department, MN Department of Natural Resources, IL Nature Preserves Commission, IL Department of Natural Resources, US Fish and Wildlife Services, US Forest Service, US Bureau of Land Management, US Army Corps of Engineers, and US National Park Service. Funding was provided by the Botanical Society of America, American Society of Plant Taxonomists, American Philosophical Society Lewis and Clark Grant, Friends of Nachusa Grasslands, Northwestern University Program in Plant Biology and Conservation, and an NSF Graduate Research Fellowship award (DGE-1842165 to K.E.W.). Undergraduate students were supported by an NSF Research Experience for Undergraduates site award to J.B.F. (DBI-461007 and DBI-1757800). Additional support was provided by the Negaunee Institute for Plant Conservation and Action at the Chicago Botanic Garden. The authors declare no conflict of interest.

## Author Contributions

K.E.W., J.B.F., and K.A.S. conceived of and planned the study. K.E.W. executed data collection with assistance from J.B.F., K.A.S. and others. K.E.W. conducted statistical analyses and wrote the manuscript with input from J.B.F., K.A.S. All authors contributed to the drafts and approved the final publication.

## Data Availability Statement

R code and data used in analyses are deposited on K. Wenzell’s GitHub (https://github.com/KWenzell).

