## Supplemental Materials for "Pollinator mosaics mirror floral trait divergence within and between species of *Castilleja*"

**FIGURES**

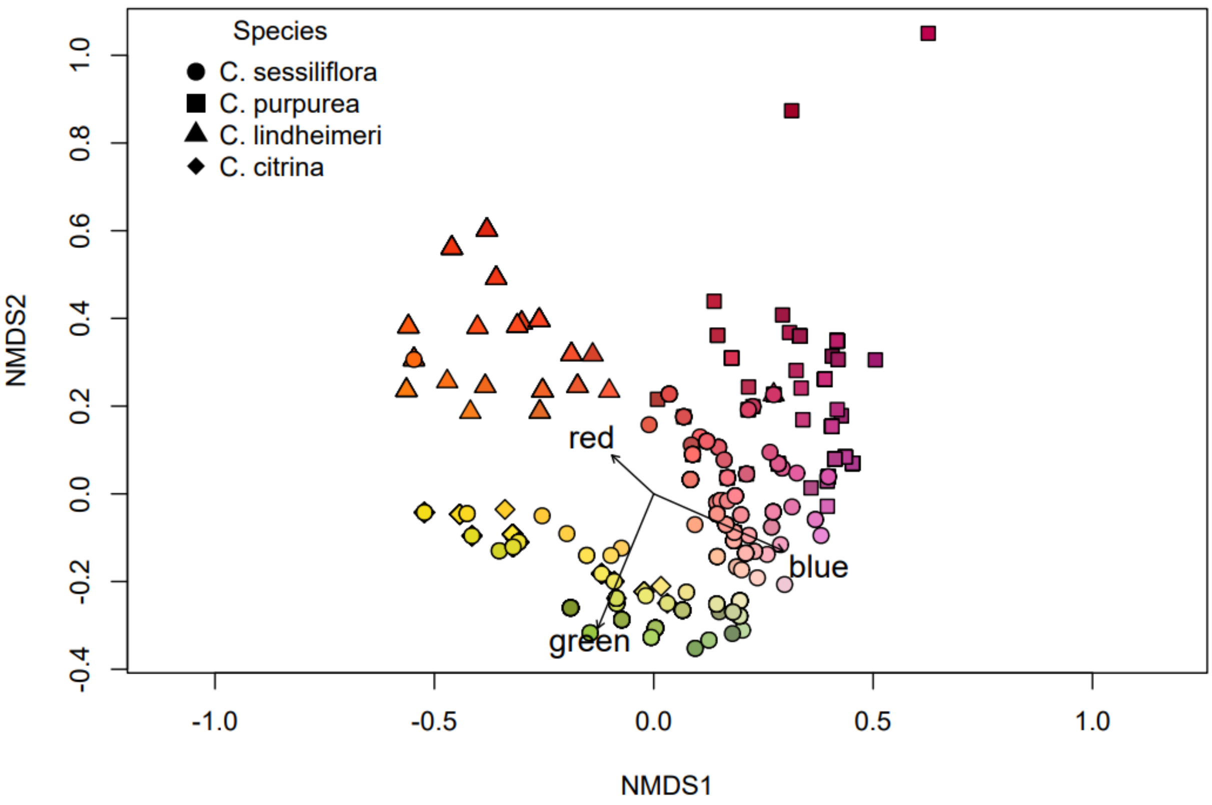


**Figure S1**. NMDS ordination of Red, Green, Blue color values of measured individuals (N=684), showing recorded inflorescence color (RGB values). Axes NMDS1 and NMDS2 reflect color variation and are used as proxies of such in analyses. Note that individual points may overlap when matched to the same color code/ RGB values.

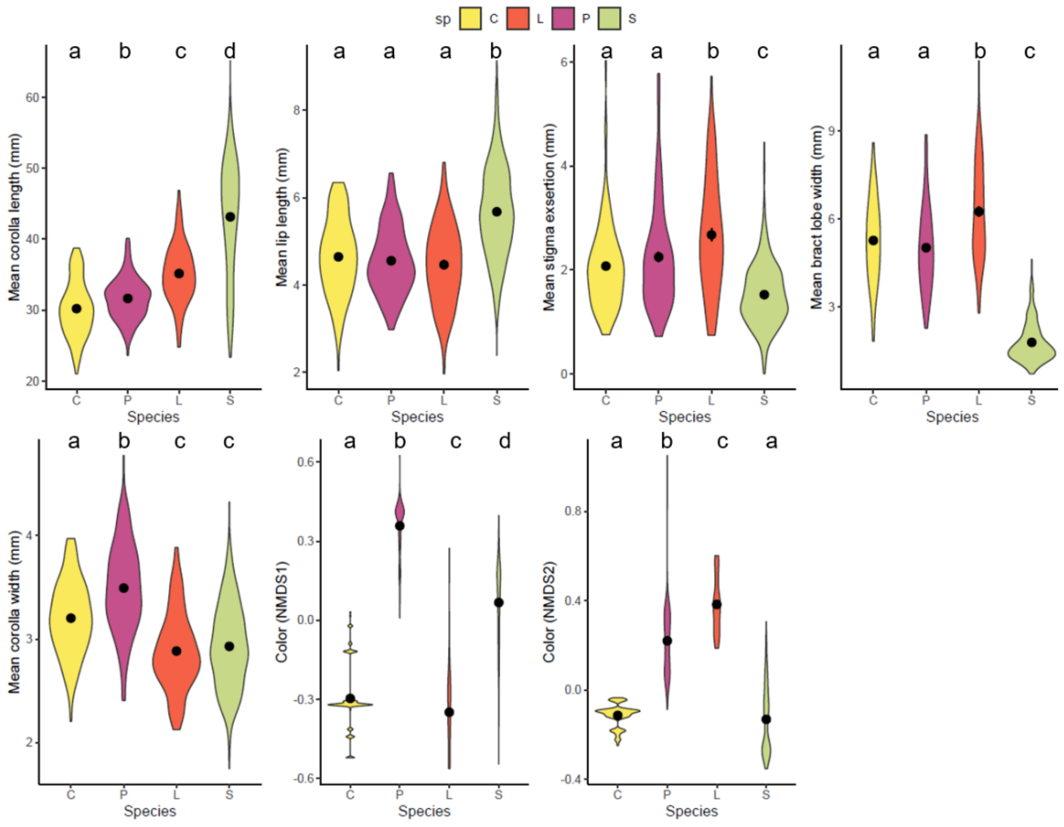


**Figure S2.** Violin plots of floral trait measurements by species (sp: C = *citrina*, L = *lindheimeri*, P = *purpurea*, S = *sessiliflora*). Central points show mean, and error bars are +/- SE. Different letters denote significant variation among species in pairwise comparison Tukey HSD tests (detailed statistics in Table S3 and S4).

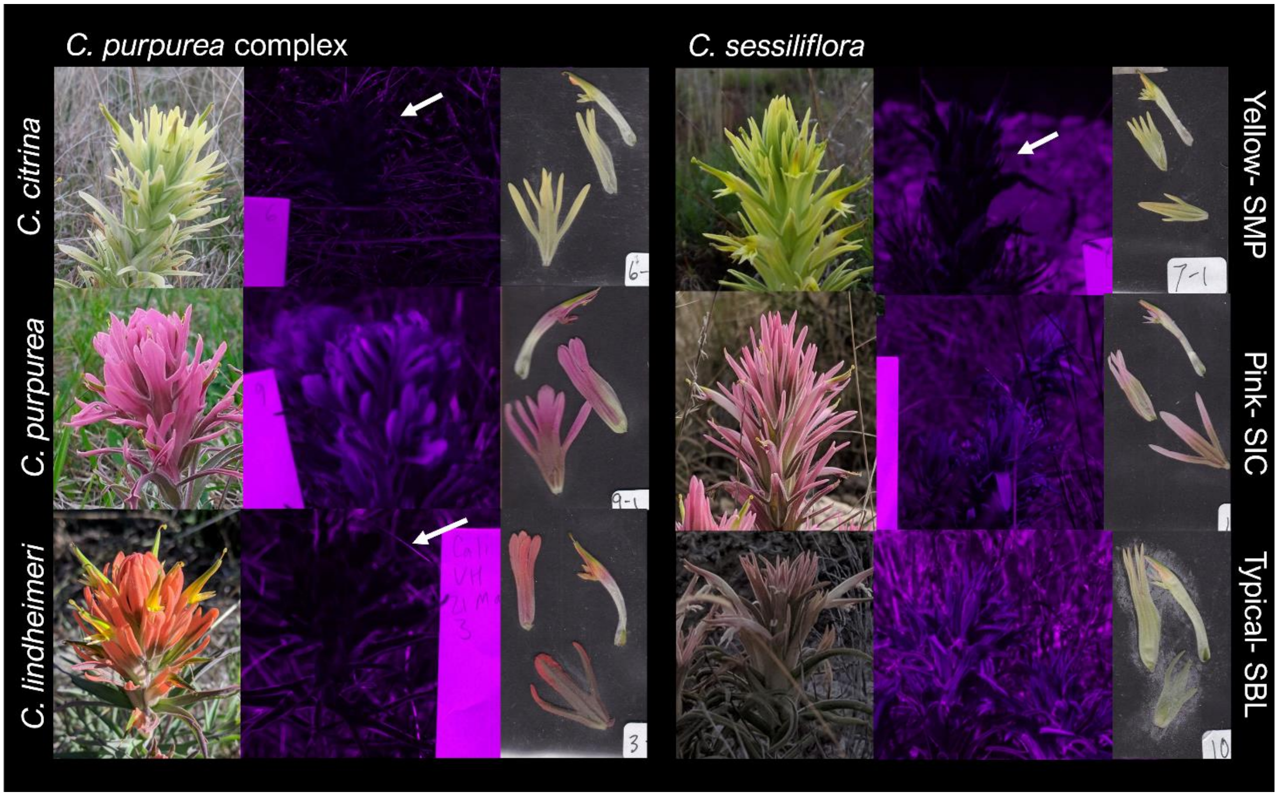


**Figure S3.** Photographs of inflorescences in human-visible wavelengths (left columns) and in UV light (center columns), along with scans of flower tissues (corolla, calyx, and bract tissues; right columns) taken from the same plants shown in UV photos. Human-visible wavelength photographs show different individuals from the same population as UV photos. Arrows point to difficult-to-see inflorescences in UV photos. Sampled populations: CMSC (*C. citrina*), PSD (*C. purpurea*, not sampled for pollinators, see Wenzell et al., 2021); LVH (*C. lindheimeri*); SMP (yellow morph *C. sessiliflora*); SIC (pink morph *C. sessiliflora*); SBL (typical morph *C. sessiliflora*). Photos not to scale.

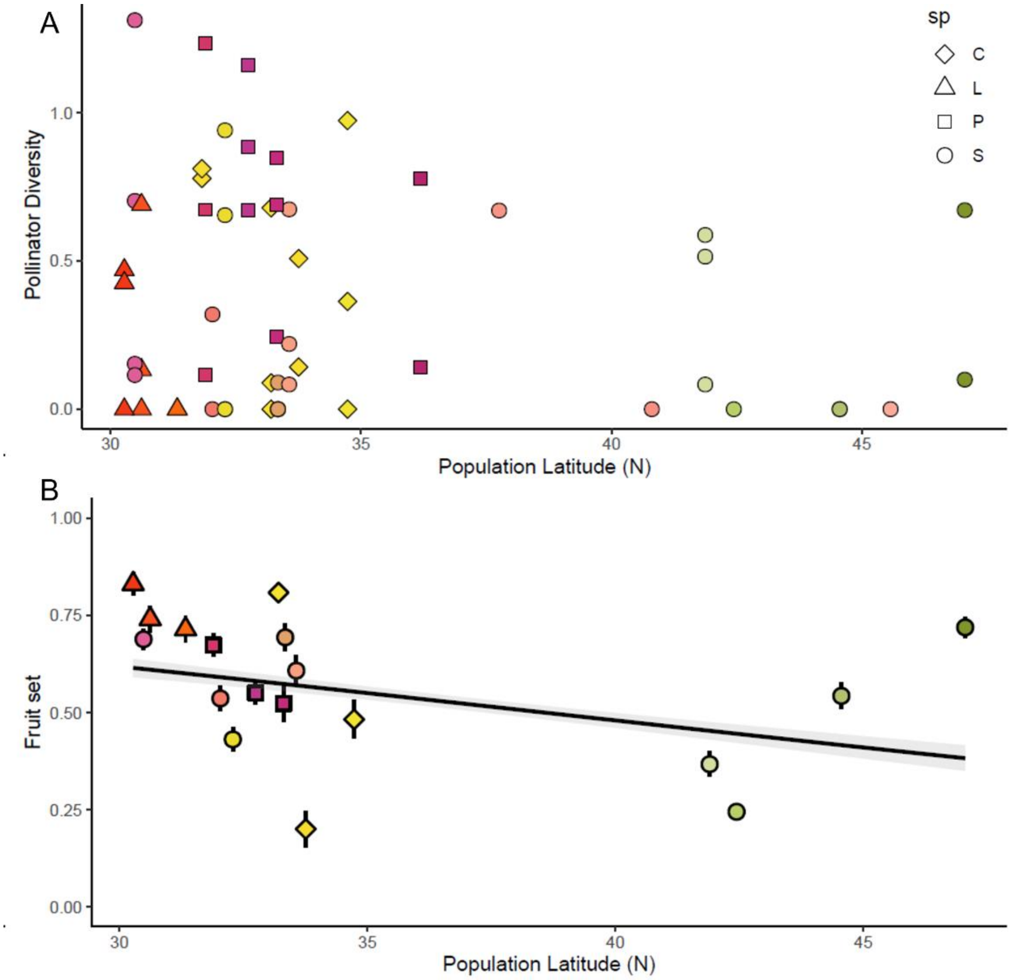


**Figure S4.** Pollinator Diversity Index (A) and population average fruit set (B) by population latitude. Species coded by shape (sp: C = *C. citrina*, L = *C. lindheimeri*, P = *C. purpurea*, S = *C. sessiliflora*), and fill color shows population median floral color, with error bars showing SE for population-average fruit set data. Solid trend lines denote a significant relationship ($p < 0.05$) based on GLMM.

### TABLES

**Table S1.** Focal populations and their location, approximate floral color (Wenzell et al., 2021, see Fig. 2), and years sampled for pollinator observations. County location is provided in place of precise coordinates for populations of conservation concern and/or in accordance with permitting regulations. Asterisks (*) denote populations sampled using the wide-view observation method, which was only utilized in 2019 (in addition to the narrow-view method used in all years). Daggers (†) denote populations and years sampled for fruit set; double daggers (‡) denote population-year datapoints included in analyses of fruit set and pollinator visitation (Fig. 6).

| Population Code | Population Name | Species | State | Latitude | Longitude | Years sampled | Floral color |
|---|---|---|---|---|---|---|---|
| CHCL | Hords Creek Lake | *C. citrina* | TX | 31.84946 | -99.58669 | 2019* | yellow |
| CLA | Lake Arrowhead | *C. citrina* | TX | 33.75562 | -98.3829 | 2018, 2019*†‡ | yellow |
| CMSC | Mt Scott Canyon | *C. citrina* | OK | 34.7308 | -98.56952 | 2018, 2019*†‡ | yellow |
| CQL | Quarry Lake | *C. citrina* | TX | 33.20647 | -98.15727 | 2018, 2019*†‡ | yellow |
| LMN | Mother Neff | *C. lindheimeri* | TX | 31.33262 | -97.46821 | 2019*†‡ | red-orange |
| LRR | Rimrock | *C. lindheimeri* | TX | 30.61922 | -98.07169 | 2018, 2019*†‡ | red-orange |
| LVH | Vireo Hill | *C. lindheimeri* | TX | 30.276778 | -97.8942907 | 2018, 2019*†‡ | red-orange |
| PCM | Clymer Meadow | *C. purpurea* | TX | 33.31087 | -96.24411 | 2018, 2019*†‡ | purple |

| | | | | | | | |
|---|---|---|---|---|---|---|---|
| PMT | Meridian Triangle | *C. purpurea* | TX | 31.88892 | -97.69912 | 2018, 2019*†‡ | purple |
| PTMS | Tulsa Mulch Site | *C. purpurea* | OK | 36.1815 | -95.8168383 | 2019* | purple |
| PTH | Tandy Hills | *C. purpurea* | TX | 32.74596 | -97.27286 | 2018, 2019*†‡ | purple |
| SFP | Felton Prairie | *C. sessiliflora* | MN | 47.05174 | -96.42735 | 2018†‡, 2019* | white-green |
| SILB | Illinois Beach | *C. sessiliflora* | IL | Lake County | - | 2017†‡, 2018†‡ | white-green |
| SNG | Nachusa Grasslands | *C. sessiliflora* | IL | Lee and Ogle Counties | - | 2017† (floral trait and fruit set data only), 2018†‡, 2019* | white-green |
| SRS | Red Shale | *C. sessiliflora* | MT | 45.57049 | -106.14787 | 2017 | pale pink |
| SSC | Spring Creek | *C. sessiliflora* | MN | 44.55531 | -92.59422 | 2017, 2018†‡ | white-green |
| SIC | Indepen-dence Creek | *C. sessiliflora* | TX | 30.486035 | -101.788252 | 2017, 2018, 2019*†‡ | pink (divergent morph) |
| SMP | Maddin Prairie | *C. sessiliflora* | TX | Mitchell County | - | 2017, 2018, 2019*†‡ | yellow (divergent morph) |
| SYH | Yeso Hills | *C. sessiliflora* | NM | 32.03468 | -104.4479 | 2018, 2019*†‡ | pale pink |
| SBL | Bottomless Lakes | *C. sessiliflora* | NM | 33.34243 | -104.33299 | 2017, 2018, 2019*†‡ | pale pink |
| SCL | Canyon Lake | *C. sessiliflora* | TX | 33.5655 | -101.80032 | 2018, 2019*†‡ | pale pink |

| | | | | | | | |
|---|---|---|---|---|---|---|---|
| SDC | David Canyon Road | *C. sessiliflora* | CO | 37.75643 | -103.59391 | 2017 | pale pink |
| SPB | Pawnee Buttes | *C. sessiliflora* | CO | 40.80505 | -104.01325 | 2017 | pale pink |

**Table S2.** Floral visitation data by species, population (Pop.), observation method (Dataset), and year. Data are presented as counts (number of visits), visitation rates (Rate; number of visits/ flower/ hour) and proportion of visits (Prop.; visits by each pollinator group divided by total visits) for each population-year-dataset datapoint by pollinator functional group, along with total number of visits, number of available flowers recorded (Total flrs), and Pollinator Diversity Index (Div. Index; log-transformed inverse Simpson's Diversity Index). Note that number of available flowers is an estimated value for Wide-view dataset observations (number of flowering stems in wide view multiplied by average number of flowers per stem recorded on narrow-view plants; see text for details). (See attached Excel file.)

**Table S3.** ANOVA results of floral traits by species and population nested within species. N = 684. Residual degrees of freedom for all models = 661.

| Trait | Variable | Degrees of Freedom | F | p |
|---|---|---|---|---|
| **Corolla length** | Species | 3 | 559.98 | **<0.0001** |
| | Sp:pop | 19 | 74.08 | **<0.0001** |
| **Corolla width** | Species | 3 | 97.27 | **<0.0001** |
| | Sp:pop | 19 | 14.74 | **<0.0001** |
| **Lip length** | Species | 3 | 121.03 | **<0.0001** |
| | Sp:pop | 19 | 21.15 | **<0.0001** |
| **Stigma** | Species | 3 | 68.329 | **<0.0001** |
| | Sp:pop | 19 | 4.558 | **<0.0001** |
| **Bract Lobe Width** | Species | 3 | 790.94 | **<0.0001** |
| | Sp:pop | 19 | 12.94 | **<0.0001** |
| **NMDS 1 (RGB)** | Species | 3 | 1232.93 | **<0.0001** |
| | Sp:pop | 19 | 42.18 | **<0.0001** |
| **NMDS 2 (RGB)** | Species | 3 | 736.49 | **<0.0001** |
| | Sp:pop | 19 | 20.94 | **<0.0001** |

**Table S4**. Tukey HSD pairwise comparisons of floral traits by species. Contrasts show pairwise comparisons between pairs of species (S = *C. sessiliflora*, C = *C. citrina*, L = *C. lindheimeri*, P = *C. purpurea*).

| | **Species** | **Mean** | **95% CI** | | **Adjusted** |
|---|---|---|---|---|---|
| **Corolla length** | **contrast** | **difference** | **Lower** | **Upper** | **p** |
| | **L-C** | **4.95** | **3.63** | **6.26** | **<0.0001** |
| | **P-C** | **1.43** | **0.22** | **2.65** | **0.013** |
| | **S-C** | **12.95** | **11.96** | **13.94** | **<0.0001** |
| | **P-L** | **-3.51** | **-4.83** | **-2.20** | **<0.0001** |
| | **S-L** | **8.01** | **6.89** | **9.12** | **<0.0001** |
| | **S-P** | **11.52** | **10.53** | **12.51** | **<0.0001** |
| **Corolla width** | **L-C** | **-0.32** | **-0.44** | **-0.20** | **<0.0001** |
| | **P-C** | **0.29** | **0.18** | **0.40** | **<0.0001** |
| | **S-C** | **-0.27** | **-0.36** | **-0.18** | **<0.0001** |
| | **P-L** | **0.61** | **0.49** | **0.73** | **<0.0001** |
| | S-L | 0.05 | -0.06 | 0.15 | 0.665 |
| | **S-P** | **-0.56** | **-0.66** | **-0.47** | **<0.0001** |
| **Lip length** | L-C | -0.18 | -0.46 | 0.09 | 0.328 |
| | P-C | -0.09 | -0.34 | 0.17 | 0.804 |
| | **S-C** | **1.03** | **0.82** | **1.24** | **<0.0001** |
| | P-L | 0.09 | -0.18 | 0.37 | 0.825 |
| | **S-L** | **1.21** | **0.98** | **1.45** | **<0.0001** |
| | **S-P** | **1.12** | **0.91** | **1.33** | **<0.0001** |
| **Stigma** | **L-C** | **0.60** | **0.32** | **0.88** | **<0.0001** |
| | P-C | 0.17 | -0.08 | 0.43 | 0.299 |
| | **S-C** | **-0.55** | **-0.76** | **-0.34** | **<0.0001** |
| | **P-L** | **-0.43** | **-0.70** | **-0.15** | **<0.001** |
| | **S-L** | **-1.15** | **-1.39** | **-0.92** | **<0.0001** |
| | **S-P** | **-0.72** | **-0.93** | **-0.51** | **<0.0001** |
| **Bract lobe width** | **L-C** | **0.98** | **0.62** | **1.34** | **<0.0001** |
| | P-C | -0.25 | -0.58 | 0.09 | 0.225 |
| | **S-C** | **-3.48** | **-3.75** | **-3.20** | **<0.0001** |

| | | | | | |
|---|---|---|---|---|---|
| | **P-L** | **-1.23** | **-1.59** | **-0.87** | **<0.0001** |
| | **S-L** | **-4.46** | **-4.76** | **-4.15** | **<0.0001** |
| | **S-P** | **-3.23** | **-3.50** | **-2.96** | **<0.0001** |
| **NMDS1 (RGB)** | **L-C** | **-0.05** | **-0.09** | **-0.02** | **0.002** |
| | **P-C** | **0.66** | **0.62** | **0.69** | **<0.0001** |
| | **S-C** | **0.36** | **0.34** | **0.39** | **<0.0001** |
| | **P-L** | **0.71** | **0.67** | **0.74** | **<0.0001** |
| | **S-L** | **0.42** | **0.39** | **0.45** | **<0.0001** |
| | **S-P** | **-0.29** | **-0.32** | **-0.26** | **<0.0001** |
| **NMDS2 (RGB)** | **L-C** | **0.50** | **0.46** | **0.54** | **<0.0001** |
| | **P-C** | **0.33** | **0.30** | **0.37** | **<0.0001** |
| | S-C | -0.02 | -0.05 | 0.01 | 0.422 |
| | **P-L** | **-0.16** | **-0.20** | **-0.12** | **<0.0001** |
| | **S-L** | **-0.51** | **-0.55** | **-0.48** | **<0.0001** |
| | **S-P** | **-0.35** | **-0.38** | **-0.32** | **<0.0001** |

**Table S5.** Variation in floral visitors by species. Model output for GLMMs of proportion and count data assessing visitation by each pollinator functional group to species, with dataset type as fixed effect and year as random effect. Proportion data models were run with a betabinomial error distribution and count models used a negative binomial distribution. Note that the count model for hummingbirds was run excluding *C. sessiliflora* due to model convergence issues expected from *C. sessiliflora* having zero hummingbird visits and accounting for about half the datapoints in the model. Residual degrees of freedom for all models = 54 (except for hummingbird count model, where residual Df = 24).

| **Proportion** | **Variable** | **$\chi^2$** | **Df** | **p** |
|---|---|---|---|---|
| Hawkmoth | Species | 2.48 | 3 | 0.479 |
| | dataset | 0.43 | 1 | 0.511 |
| Small-med bee | Species | 7.05 | 3 | 0.070 |
| | **dataset** | 16.68 | 1 | **<0.0001** |
| Bumblebee | Species | 4.06 | 3 | 0.255 |
| | **dataset** | 6.41 | 1 | **0.011** |
| Hummingbird | **Species** | 16.35 | 3 | **0.001** |
| | **dataset** | 6.25 | 1 | **0.012** |
| Butterfly | **Species** | 12.49 | 3 | **0.006** |
| | **dataset** | 6.55 | 1 | **0.011** |
| Other | Species | 5.87 | 3 | 0.118 |
| | dataset | 0.53 | 1 | 0.467 |

| **Count** | **Variable** | **$\chi^2$** | **Df** | **p** |
|---|---|---|---|---|
| Hawkmoth | Species | 6.33 | 3 | 0.097 |
| | dataset | 0.68 | 1 | 0.409 |
| Small-med bee | **Species** | 8.09 | 3 | **0.044** |
| | **dataset** | 6.85 | 1 | **0.009** |

| | | | | |
|---|---|---|---|---|
| Bumblebee | Species | 2.57 | 3 | 0.463 |
| | **dataset** | 4.25 | 1 | **0.039** |
| Hummingbird | **Species** | 20.99 | 2,24 | **<0.0001** |
| | **dataset** | 17.04 | 1,24 | **<0.0001** |
| Butterfly | **Species** | 13.86 | 3 | **0.003** |
| | **dataset** | 5.94 | 1 | **0.015** |
| Other | Species | 7.76 | 3 | 0.051 |
| | dataset | 0.01 | 1 | 0.906 |

**Table S6.** Pairwise multiple comparisons of proportion and number of visits (count) by pollinator group among species, presented only for pollinator functional groups with significant variation among species in our GLMM. Contrasts show pairwise comparisons between pairs of species (S = *C. sessiliflora*, C = *C. citrina*, L = *C. lindheimeri*, P = *C. purpurea*) averaged over year and dataset. Note that p = 1.0 results from comparisons to groups with all zeroes for the response variable (resulting in zero variance and high SE). Contrasts in bold are significantly different (p < 0.05). Note that the count model for hummingbirds was run excluding *C. sessiliflora* due to model convergence issues due to large number of zeroes.

**Hummingbirds: proportion**

| contrast | Estimate | SE | Df | t ratio | p |
|---|---|---|---|---|---|
| **C-L** | **-3.84** | **1.07** | **54** | **-3.58** | **0.004** |
| C-P | -0.56 | 0.94 | 54 | -0.59 | 0.934 |
| C-S | 20.70 | 11300.00 | 54 | 0.00 | 1.000 |
| **L-P** | **3.28** | **0.92** | **54** | **3.56** | **0.004** |
| L-S | 24.54 | 11300.00 | 54 | 0.00 | 1.000 |
| P-S | 21.26 | 11300.00 | 54 | 0.00 | 1.000 |

**Butterflies: proportion**

| contrast | Estimate | SE | Df | t ratio | p |
|---|---|---|---|---|---|
| C-L | 0.14 | 0.93 | 54 | 0.15 | 0.999 |

| | | | | | |
|---|---|---|---|---|---|
| C-P | -1.14 | 0.70 | 54 | -1.62 | 0.375 |
| C-S | 1.20 | 0.84 | 54 | 1.43 | 0.486 |
| L-P | -1.28 | 0.81 | 54 | -1.59 | 0.393 |
| L-S | 1.06 | 0.93 | 54 | 1.14 | 0.667 |
| **P-S** | **2.34** | **0.69** | **54** | **3.39** | **0.007** |

**Small/med bees: proportion**

| **contrast** | **Estimate** | **SE** | **Df** | **t ratio** | **p** |
|---|---|---|---|---|---|
| C-L | 1.58 | 0.80 | 54 | 1.97 | 0.213 |
| C-P | 0.06 | 0.63 | 54 | 0.10 | 1.000 |
| C-S | -0.30 | 0.55 | 54 | -0.54 | 0.949 |
| L-P | -1.52 | 0.77 | 54 | -1.98 | 0.208 |
| L-S | -1.88 | 0.71 | 54 | -2.65 | 0.050 |
| P-S | -0.36 | 0.50 | 54 | -0.72 | 0.889 |

**Hummingbirds: count (*C. purpurea* complex only)**

| **contrast** | **Estimate** | **SE** | **Df** | **t ratio** | **p** |
|---|---|---|---|---|---|
| **C-L** | **-2.76** | **0.76** | **24** | **-3.66** | **0.004** |
| C-P | -0.83 | 0.84 | 24 | -0.98 | 0.596 |
| **L-P** | **1.94** | **0.56** | **24** | **3.46** | **0.006** |

**Butterflies: count**

| **contrast** | **Estimate** | **SE** | **Df** | **t ratio** | **p** |
|---|---|---|---|---|---|
| C-L | -0.16 | 0.91 | 54 | -0.18 | 0.998 |
| C-P | -1.36 | 0.67 | 54 | -2.03 | 0.190 |
| C-S | 0.97 | 0.82 | 54 | 1.19 | 0.637 |
| L-P | -1.19 | 0.77 | 54 | -1.54 | 0.419 |
| L-S | 1.13 | 0.91 | 54 | 1.25 | 0.598 |
| **P-S** | **2.33** | **0.67** | **54** | **3.47** | **0.006** |

| **Small/med bees: count** | | | | | |
|---|---|---|---|---|---|
| **contrast** | **Estimate** | **SE** | **Df** | **t ratio** | **p** |
| C-L | 0.97 | 0.68 | 54 | 1.42 | 0.495 |
| C-P | -0.78 | 0.47 | 54 | -1.66 | 0.352 |
| C-S | -0.27 | 0.45 | 54 | -0.60 | 0.931 |
| **L-P** | **-1.74** | **0.65** | **54** | **-2.69** | **0.046** |
| L-S | -1.24 | 0.63 | 54 | -1.97 | 0.211 |
| P-S | 0.51 | 0.38 | 54 | 1.33 | 0.547 |

**Table S7.** Latitudinal variation in floral visitors within *C. sessiliflora*. Model output for GLMMs of proportion of visitation against population latitude for each pollinator group, with dataset type as fixed effect. Models used a betabinomial distribution, residual degrees of freedom for all models = 26, N =31.

| **Group** | **Variable** | **$\chi^2$** | **Df** | **p** |
|---|---|---|---|---|
| Hawkmoth | **Latitude** | **11.102** | **1** | **0.001** |
| | dataset | 0.000 | 1 | 0.983 |
| Small-med bee | **Latitude** | **14.899** | **1** | **< 0.001** |
| | **dataset** | **12.398** | **1** | **< 0.001** |
| Bumblebee | Latitude | 1.698 | 1 | 0.193 |
| | **dataset** | **7.430** | **1** | **0.006** |
| Other | Latitude | 0.383 | 1 | 0.536 |
| | dataset | 0.464 | 1 | 0.496 |

**Table S8.** Pairwise multiple comparisons of Pollinator Diversity Index among species, based on GLMM. Contrasts show pairwise comparisons between pairs of species (S = *C. sessiliflora*, C = *C. citrina*, L = *C. lindheimeri*, P = *C. purpurea*) averaged over dataset.

**Pollinator Diversity by Species**

| contrast | Estimate | SE | Df | t ratio | p |
|---|---|---|---|---|---|
| C-L | 0.22 | 0.16 | 50 | 1.37 | 0.522 |
| C-P | -0.24 | 0.15 | 50 | -1.61 | 0.384 |
| C-S | 0.15 | 0.13 | 50 | 1.19 | 0.637 |
| **L-P** | **-0.46** | **0.16** | **50** | **-2.92** | **0.026** |
| L-S | -0.07 | 0.14 | 50 | -0.53 | 0.952 |
| **P-S** | **0.39** | **0.12** | **50** | **3.19** | **0.013** |

**Table S9.** GLMM results of population average fruit set in relation to proportion visitation by each pollinator functional group, pollinator diversity, and total number of visits. Dataset type was included as a fixed effect. Coefficient estimate (Estimate, in logit scale), standard error (SE), z-value (z) and p-value (p) are presented for each variable, along with $\chi^2$ and degrees of freedom (Df, with residual Df). P < 0.05 and associated variables are in bold. Note no hummingbird visits were recorded to *C. sessiliflora*.

| Focal Group | Variable | Estimate | SE | z | $\chi^2$ | Df | p |
|---|---|---|---|---|---|---|---|
| *C. sessiliflora* | **Hawkmoth** | **0.744** | **0.343** | **2.169** | **4.706** | **1,11** | **0.030** |
| | dataset | 0.263 | 0.341 | 0.772 | 0.595 | 1,11 | 0.440 |
| *C. purpurea* complex | Hawkmoth | -0.820 | 0.650 | -1.263 | 1.594 | 1,13 | 0.207 |
| | dataset | -0.071 | 0.314 | -0.226 | 0.051 | 1,13 | 0.821 |
| *C. sessiliflora* | Hummingbird | NA | | | | | |
| | dataset | NA | | | | | |
| *C. purpurea* complex | **Hummingbird** | **1.058** | **0.363** | **2.916** | **8.504** | **1,13** | **0.004** |
| | dataset | -0.149 | 0.291 | -0.511 | 0.261 | 1,13 | 0.609 |
| *C. sessiliflora* | Small/medium bee | -0.446 | 0.439 | -1.015 | 1.031 | 1,11 | 0.310 |

| | | | | | | | |
|---|---|---|---|---|---|---|---|
| | dataset | 0.208 | 0.410 | 0.509 | 0.259 | 1,11 | 0.611 |
| *C. purpurea* complex | Small/medium bee | -0.025 | 0.850 | -0.029 | 0.001 | 1,13 | 0.977 |
| | dataset | -0.008 | 0.421 | -0.019 | 0.000 | 1,13 | 0.985 |
| *C. sessiliflora* | Large bee | -0.788 | 0.724 | -1.088 | 1.184 | 1,11 | 0.276 |
| | dataset | 0.595 | 0.405 | 1.468 | 2.156 | 1,11 | 0.142 |
| *C. purpurea* complex | Large bee | 0.911 | 0.863 | 1.055 | 1.113 | 1,13 | 0.291 |
| | dataset | -0.132 | 0.322 | -0.409 | 0.167 | 1,13 | 0.683 |
| *C. sessiliflora* | Butterfly | 13.592 | 12.942 | 1.050 | 1.103 | 1,11 | 0.294 |
| | dataset | 0.292 | 0.371 | 0.787 | 0.620 | 1,11 | 0.431 |
| *C. purpurea* complex | Butterfly | -0.472 | 1.162 | -0.407 | 0.166 | 1,13 | 0.684 |
| | dataset | 0.058 | 0.343 | 0.168 | 0.028 | 1,13 | 0.866 |
| *C. sessiliflora* | Other visitors | 3.074 | 8.986 | 0.342 | 0.117 | 1,11 | 0.732 |
| | dataset | 0.428 | 0.381 | 1.126 | 1.267 | 1,11 | 0.260 |
| *C. purpurea* complex | Other visitors | 0.563 | 1.242 | 0.454 | 0.206 | 1,13 | 0.650 |
| | dataset | 0.043 | 0.324 | 0.133 | 0.018 | 1,13 | 0.894 |
| *C. sessiliflora* | Pollinator Diversity | 1.194 | 0.877 | 1.362 | 1.856 | 1,8 | 0.173 |
| | dataset | 0.407 | 0.330 | 1.234 | 1.523 | 1,8 | 0.217 |
| *C. purpurea* complex | Pollinator Diversity | -0.207 | 0.455 | -0.454 | 0.206 | 1,12 | 0.650 |
| | dataset | 0.203 | 0.290 | 0.702 | 0.493 | 1,12 | 0.483 |
| *C. sessiliflora* | Total visits | 0.002 | 0.006 | 0.339 | 0.115 | 1,11 | 0.735 |
| | dataset | 0.351 | 0.393 | 0.893 | 0.798 | 1,11 | 0.372 |
| *C. purpurea* complex | Total visits | 0.002 | 0.003 | 0.715 | 0.511 | 1,13 | 0.475 |
| | dataset | -0.226 | 0.444 | -0.509 | 0.259 | 1,13 | 0.611 |

**Supplemental Methods:**

Details on methods for classifying pollinator functional groups and recording floral visits:

Pollinator functional groups included: hawkmoths (exclusively *Hyles lineata*, Sphingidae, based on individuals that could be identified on the wing or in photographs), hummingbirds (Trochilidae), bumblebees (*Bombus* sp.), other large bees (approximately 2.5 cm or larger in body length, such as carpenter bees), medium bees (approximately 1.5 cm in length), small bees (approximately <1 cm in length, including sweat bees such as *Lassioglossum* sp., and other pollen-collecting bees), bee flies (Bombyliidae), large butterflies (approximately > 4 cm in wing height, including swallowtail butterflies such as *Battus philenor* and others), small butterflies (approximately < 3 cm), non-Sphingid moths, and flies.

Methods for count data models of floral visitation among species:

Models of visit counts used number of floral visits per pollinator group as the response term and plant species as the predictor, with a negative binomial error distribution. The model of hummingbird visit count data was run on only the species of *C. purpurea* complex (due to zero hummingbird visits recorded to *C. sessiliflora*). To assess potential overdispersion in the data, we ran models using both a Poisson and a negative binomial error distribution and used package DHARMa (Hartig, 2016) to examine residuals as described for proportion models. The count models with Poisson distributions showed greater evidence of overdispersion than those using negative binomial distributions, and therefore negative binomial error distributions were chosen for count models.
