## Supplemental Table S2 for "Pollinator mosaics mirror floral trait divergence within and between species of *Castilleja*"

**Table S2.** Floral visitation data by species, population (Pop.), observation method (Dataset), and year. Data are presented as counts (number of visits), visitation rates (Rate; number of visits/ flower/ hour) and proportion of visits (Prop.; visits by each pollinator group divided by total visits) for each population-year-dataset datapoint by pollinator functional group, along with total number of visits, number of available flowers recorded (Total flrs), and Pollinator Diversity Index (Div. Index; log-transformed inverse Simpson's Diversity Index). Note that number of available flowers is an estimated value for Wide-view dataset observations (number of flowering stems in wide view multiplied by average number of flowers per stem recorded on narrow-view plants; see text for details).

| ***C. purpurea* complex** | | | **Small/ medium bees** | | | **Large/ Bumblebees** | | | **Hawkmoths** | | |
|---|---|---|---|---|---|---|---|---|---|---|---|
| Pop. | Dataset | Year | Count | Rate | Prop. | Count | Rate | Prop. | Count | Rate | Prop. |
| *C. citrina* | | | | | | | | | | | |
| CHCL | Narrow view | 2019 | 0 | 0 | 0 | 8 | 0.0078 | 0.57143 | 5 | 0.005 | 0.35714 |
| | Wide view | 2019 | 0 | 0 | 0 | 163 | 0.0141 | 0.62214 | 13 | 0.001 | 0.04962 |
| CLA | Narrow view | 2018 | 3 | 0 | 0.1875 | 12 | 0.0044 | 0.75 | 1 | 4E-04 | 0.0625 |
| | | 2019 | 13 | 0.01 | 0.9286 | 0 | 0 | 0 | 0 | 0 | 0 |
| | Wide view | 2019 | 0 | 0 | 0 | 0 | 0 | 0 | 0 | 0 | 0 |
| CMSC | Narrow view | 2018 | 15 | 0.01 | 0.4054 | 16 | 0.0141 | 0.43243 | 6 | 0.005 | 0.16216 |
| | | 2019 | 0 | 0 | 0 | 0 | 0 | 0 | 11 | 0.024 | 1 |
| | Wide view | 2019 | 3 | 0 | 0.1875 | 0 | 0 | 0 | 13 | 0.008 | 0.8125 |
| CQL | Narrow view | 2018 | 43 | 0.03 | 0.9556 | 2 | 0.0012 | 0.04444 | 0 | 0 | 0 |
| | | 2019 | 1 | 0 | 1 | 0 | 0 | 0 | 0 | 0 | 0 |
| | Wide view | 2019 | 7 | 0 | 0.068 | 67 | 0.0064 | 0.65049 | 0 | 0 | 0 |
| *C. lindheimeri* | | | | | | | | | | | |
| LMN | Narrow view | 2019 | 0 | 0 | 0 | 0 | 0 | 0 | 0 | 0 | 0 |
| | Wide view | 2019 | 0 | 0 | 0 | 0 | 0 | 0 | 0 | 0 | 0 |
| LRR | Narrow view | 2018 | 2 | 0 | 0.0667 | 0 | 0 | 0 | 0 | 0 | 0 |
| | | 2019 | 0 | 0 | 0 | 0 | 0 | 0 | 0 | 0 | 0 |
| | Wide view | 2019 | 0 | 0 | 0 | 0 | 0 | 0 | 0 | 0 | 0 |
| LVH | Narrow view | 2018 | 0 | 0 | 0 | 0 | 0 | 0 | 17 | 0.025 | 1 |
| | | 2019 | 1 | 0 | 0.25 | 0 | 0 | 0 | 0 | 0 | 0 |
| | Wide view | 2019 | 1 | 0 | 0.0182 | 0 | 0 | 0 | 3 | 9E-04 | 0.05455 |
| *C. purpurea* | | | | | | | | | | | |
| PCM | Narrow view | 2018 | 8 | 0.01 | 0.08 | 0 | 0 | 0 | 88 | 0.095 | 0.88 |
| | | 2019 | 6 | 0.01 | 0.2857 | 0 | 0 | 0 | 12 | 0.017 | 0.57143 |
| | Wide view | 2019 | 0 | 0 | 0 | 59 | 0.0134 | 0.66292 | 21 | 0.005 | 0.23596 |
| PMT | Narrow view | 2018 | 23 | 0.02 | 0.3382 | 21 | 0.0152 | 0.30882 | 19 | 0.014 | 0.27941 |
| | | 2019 | 3 | 0 | 0.4286 | 0 | 0 | 0 | 0 | 0 | 0 |
| | Wide view | 2019 | 0 | 0 | 0 | 0 | 0 | 0 | 0 | 0 | 0 |
| PTH | Narrow view | 2018 | 101 | 0.02 | 0.3556 | 1 | 0.0002 | 0.00352 | 176 | 0.035 | 0.61972 |
| | | 2019 | 29 | 0.01 | 0.3537 | 0 | 0 | 0 | 43 | 0.013 | 0.52439 |
| | Wide view | 2019 | 8 | 0 | 0.0406 | 4 | 7E-05 | 0.0203 | 48 | 9E-04 | 0.24365 |
| PTMS | Narrow view | 2019 | 67 | 0.02 | 0.9306 | 0 | 0 | 0 | 2 | 7E-04 | 0.02778 |
| | Wide view | 2019 | 3 | 0 | 0.3 | 0 | 0 | 0 | 0 | 0 | 0 |

***C. sessiliflora***

| | | | Small/ medium bees | | | Large/ Bumblebees | | | Hawkmoths | | |
|---|---|---|---|---|---|---|---|---|---|---|---|
| Pop. | Dataset | Year | Count | Rate | Prop. | Count | Rate | Prop. | Count | Rate | Prop. |
| SBL | Narrow view | 2017 | 5 | 0.02 | 0.045 | 0 | 0 | 0 | 106 | 0.425 | 0.95495 |
| | | 2018 | 0 | 0 | 0 | 0 | 0 | 0 | 69 | 0.205 | 1 |
| | | 2019 | 0 | 0 | 0 | 0 | 0 | 0 | 4 | 0.007 | 1 |
| | Wide view | 2019 | 0 | 0 | 0 | 0 | 0 | 0 | 29 | 0.06 | 1 |
| SCL | Narrow view | 2018 | 50 | 0.05 | 0.431 | 0 | 0 | 0 | 66 | 0.067 | 0.56897 |
| | | 2019 | 2 | 0 | 0.0417 | 0 | 0 | 0 | 46 | 0.086 | 0.95833 |
| | Wide view | 2019 | 0 | 0 | 0 | 2 | 0.002 | 0.11111 | 16 | 0.016 | 0.88889 |
| SDCR | Narrow view | 2017 | 78 | 0.18 | 0.4239 | 0 | 0 | 0 | 106 | 0.241 | 0.57609 |
| SFP | Narrow view | 2018 | 0 | 0 | 0 | 0 | 0 | 0 | 0 | 0 | 0 |
| | | 2019 | 35 | 0.06 | 0.6481 | 16 | 0.0287 | 0.2963 | 0 | 0 | 0 |
| | Wide view | 2019 | 4 | 0 | 0.0396 | 96 | 0.0814 | 0.9505 | 0 | 0 | 0 |
| SIC | Narrow view | 2017 | 85 | 0.15 | 0.2911 | 0 | 0 | 0 | 53 | 0.095 | 0.18151 |
| | | 2018 | 12 | 0.08 | 0.9231 | 0 | 0 | 0 | 0 | 0 | 0 |
| | | 2019 | 20 | 0.03 | 0.2469 | 0 | 0 | 0 | 53 | 0.076 | 0.65432 |
| | Wide view | 2019 | 0 | 0 | 0 | 0 | 0 | 0 | 49 | 0.016 | 0.94231 |
| SILB | Narrow view | 2017 | 0 | 0 | 0 | 0 | 0 | 0 | 0 | 0 | 0 |
| | | 2018 | 10 | 0.01 | 1 | 0 | 0 | 0 | 0 | 0 | 0 |
| SMP | Narrow view | 2017 | 32 | 0.04 | 0.2991 | 5 | 0.0065 | 0.04673 | 70 | 0.091 | 0.65421 |
| | | 2018 | 10 | 0.01 | 0.303 | 17 | 0.0209 | 0.51515 | 6 | 0.007 | 0.18182 |
| | | 2019 | 0 | 0 | 0 | 0 | 0 | 0 | 1 | 4E-04 | 1 |
| | Wide view | 2019 | 0 | 0 | 0 | 105 | 0.0057 | 1 | 0 | 0 | 0 |
| SNG | Narrow view | 2018 | 23 | 0.13 | 0.9583 | 0 | 0 | 0 | 0 | 0 | 0 |
| | | 2019 | 165 | 0.18 | 0.7237 | 62 | 0.0671 | 0.27193 | 0 | 0 | 0 |
| | Wide view | 2019 | 7 | 0.01 | 0.3333 | 14 | 0.0156 | 0.66667 | 0 | 0 | 0 |
| SPB | Narrow view | 2017 | 25 | 0.03 | 1 | 0 | 0 | 0 | 0 | 0 | 0 |
| SRS | Narrow view | 2017 | 36 | 0.07 | 1 | 0 | 0 | 0 | 0 | 0 | 0 |
| SSC | Narrow view | 2017 | 28 | 0.05 | 1 | 0 | 0 | 0 | 0 | 0 | 0 |
| | | 2018 | 24 | 0.13 | 1 | 0 | 0 | 0 | 0 | 0 | 0 |
| SYH | Narrow view | 2018 | 46 | 0.3 | 0.8364 | 0 | 0 | 0 | 9 | 0.058 | 0.16364 |
| | | 2019 | 1 | 0 | 1 | 0 | 0 | 0 | 0 | 0 | 0 |
| | Wide view | 2019 | 0 | 0 | 0 | 0 | 0 | 0 | 0 | 0 | 0 |

| ***C. purpurea* complex** | | | **Hummingbirds** | | | **Butterflies** | | | **Other visitors** | | | | | |
|---|---|---|---|---|---|---|---|---|---|---|---|---|---|---|
| Pop. | Dataset | Year | Count | Rate | Prop. | Count | Rate | Prop. | Count | Rate | Prop. | Total visits | Total flrs | Div. Index |
| *C. citrina* | | | | | | | | | | | | | | |
| CHCL | Narrow | 2019 | 0 | 0 | 0 | 1 | 0 | 0.071 | 0 | 0 | 0 | 14 | 140 | 0.778 |
| | Wide | 2019 | 34 | 0 | 0.13 | 51 | 0 | 0.195 | 1 | 0 | 0 | 262 | 1578 | 0.811 |
| CLA | Narrow | 2018 | 0 | 0 | 0 | 0 | 0 | 0 | 0 | 0 | 0 | 16 | 370 | 0.508 |
| | | 2019 | 0 | 0 | 0 | 1 | 0 | 0.071 | 0 | 0 | 0 | 14 | 281 | 0.142 |
| | Wide | 2019 | 0 | 0 | 0 | 0 | 0 | 0 | 0 | 0 | 0 | 0 | 347.2 | NA |
| CMSC | Narrow | 2018 | 0 | 0 | 0 | 0 | 0 | 0 | 0 | 0 | 0 | 37 | 155 | 0.974 |
| | | 2019 | 0 | 0 | 0 | 0 | 0 | 0 | 0 | 0 | 0 | 11 | 63 | 0.000 |
| | Wide | 2019 | 0 | 0 | 0 | 0 | 0 | 0 | 0 | 0 | 0 | 16 | 214.8 | 0.363 |
| CQL | Narrow | 2018 | 0 | 0 | 0 | 0 | 0 | 0 | 0 | 0 | 0 | 45 | 221 | 0.089 |
| | | 2019 | 0 | 0 | 0 | 0 | 0 | 0 | 0 | 0 | 0 | 1 | 137 | 0.000 |
| | Wide | 2019 | 29 | 0 | 0.282 | 0 | 0 | 0 | 0 | 0 | 0 | 103 | 1436 | 0.679 |
| *C. lindheimeri* | | | | | | | | | | | | | | |
| LMN | Narrow | 2019 | 16 | 0.02 | 1 | 0 | 0 | 0 | 0 | 0 | 0 | 16 | 136 | 0.000 |
| | Wide | 2019 | 209 | 0.06 | 1 | 0 | 0 | 0 | 0 | 0 | 0 | 209 | 462.3 | 0.000 |
| LRR | Narrow | 2018 | 28 | 0.03 | 0.933 | 0 | 0 | 0 | 0 | 0 | 0 | 30 | 111 | 0.133 |
| | | 2019 | 7 | 0.01 | 1 | 0 | 0 | 0 | 0 | 0 | 0 | 7 | 96 | 0.000 |
| | Wide | 2019 | 123 | 0.01 | 0.528 | 110 | 0.01 | 0.472 | 0 | 0 | 0 | 233 | 1218 | 0.690 |
| LVH | Narrow | 2018 | 0 | 0 | 0 | 0 | 0 | 0 | 0 | 0 | 0 | 17 | 94 | 0.000 |
| | | 2019 | 3 | 0.01 | 0.75 | 0 | 0 | 0 | 0 | 0 | 0 | 4 | 70 | 0.470 |
| | Wide | 2019 | 44 | 0.01 | 0.8 | 5 | 0 | 0.091 | 2 | 0 | 0.04 | 55 | 450.6 | 0.426 |
| *C. purpurea* | | | | | | | | | | | | | | |
| PCM | Narrow | 2018 | 0 | 0 | 0 | 4 | 0 | 0.04 | 0 | 0 | 0 | 100 | 127 | 0.245 |
| | | 2019 | 0 | 0 | 0 | 3 | 0 | 0.143 | 0 | 0 | 0 | 21 | 96 | 0.847 |
| | Wide | 2019 | 2 | 0 | 0.022 | 7 | 0 | 0.079 | 0 | 0 | 0 | 89 | 602.6 | 0.689 |
| PMT | Narrow | 2018 | 3 | 0 | 0.044 | 2 | 0 | 0.029 | 0 | 0 | 0 | 68 | 188 | 1.236 |
| | | 2019 | 0 | 0 | 0 | 0 | 0 | 0 | 4 | 0 | 0.57 | 7 | 280 | 0.673 |
| | Wide | 2019 | 115 | 0.01 | 0.943 | 7 | 0 | 0.057 | 0 | 0 | 0 | 122 | 2088 | 0.114 |
| PTH | Narrow | 2018 | 0 | 0 | 0 | 0 | 0 | 0 | 6 | 0 | 0.02 | 284 | 677 | 0.671 |
| | | 2019 | 0 | 0 | 0 | 1 | 0 | 0.012 | 9 | 0 | 0.11 | 82 | 442 | 0.886 |
| | Wide | 2019 | 24 | 0 | 0.122 | 94 | 0 | 0.477 | 19 | 0 | 0.1 | 197 | 7614 | 1.161 |
| PTMS | Narrow | 2019 | 0 | 0 | 0 | 0 | 0 | 0 | 3 | 0 | 0.04 | 72 | 398 | 0.141 |
| | Wide | 2019 | 0 | 0 | 0 | 6 | 0 | 0.6 | 1 | 0 | 0.1 | 10 | 3597 | 0.777 |

*C. sessiliflora*

| Pop. | Dataset | Year | **Hummingbirds** Count | Rate | Prop. | **Butterflies** Count | Rate | Prop. | **Other visitors** Count | Rate | Prop. | Total visits | Total flrs | Div. Index |
|---|---|---|---|---|---|---|---|---|---|---|---|---|---|---|
| SBL | Narrow | 2017 | 0 | 0 | 0 | 0 | 0 | 0 | 0 | 0 | 0 | 111 | 34 | 0.090 |
| | | 2018 | 0 | 0 | 0 | 0 | 0 | 0 | 0 | 0 | 0 | 69 | 46 | 0.000 |
| | | 2019 | 0 | 0 | 0 | 0 | 0 | 0 | 0 | 0 | 0 | 4 | 77 | 0.000 |
| | Wide | 2019 | 0 | 0 | 0 | 0 | 0 | 0 | 0 | 0 | 0 | 29 | 65.45 | 0.000 |
| SCL | Narrow | 2018 | 0 | 0 | 0 | 0 | 0 | 0 | 0 | 0 | 0 | 116 | 134 | 0.674 |
| | | 2019 | 0 | 0 | 0 | 0 | 0 | 0 | 0 | 0 | 0 | 48 | 73 | 0.083 |
| | Wide | 2019 | 0 | 0 | 0 | 0 | 0 | 0 | 0 | 0 | 0 | 18 | 139.9 | 0.220 |
| SDCR | Narrow | 2017 | 0 | 0 | 0 | 0 | 0 | 0 | 0 | 0 | 0 | 184 | 60 | 0.670 |
| SFP | Narrow | 2018 | 0 | 0 | 0 | 0 | 0 | 0 | 0 | 0 | 0 | 0 | 11 | NA |
| | | 2019 | 0 | 0 | 0 | 0 | 0 | 0 | 3 | 0.01 | 0.06 | 54 | 76 | 0.671 |
| | Wide | 2019 | 0 | 0 | 0 | 0 | 0 | 0 | 1 | 0 | 0.01 | 101 | 161 | 0.100 |
| SIC | Narrow | 2017 | 0 | 0 | 0 | 100 | 0.18 | 0.342 | 54 | 0.1 | 0.18 | 292 | 76 | 1.312 |
| | | 2018 | 0 | 0 | 0 | 0 | 0 | 0 | 1 | 0.01 | 0.08 | 13 | 21 | 0.153 |
| | | 2019 | 0 | 0 | 0 | 2 | 0 | 0.025 | 6 | 0.01 | 0.07 | 81 | 95 | 0.703 |
| | Wide | 2019 | 0 | 0 | 0 | 3 | 0 | 0.058 | 0 | 0 | 0 | 52 | 418 | 0.115 |
| SILB | Narrow | 2017 | 0 | 0 | 0 | 0 | 0 | 0 | 0 | 0 | 0 | 0 | 110 | NA |
| | | 2018 | 0 | 0 | 0 | 0 | 0 | 0 | 0 | 0 | 0 | 10 | 174 | 0.000 |
| SMP | Narrow | 2017 | 0 | 0 | 0 | 0 | 0 | 0 | 0 | 0 | 0 | 107 | 105 | 0.655 |
| | | 2018 | 0 | 0 | 0 | 0 | 0 | 0 | 0 | 0 | 0 | 33 | 111 | 0.941 |
| | | 2019 | 0 | 0 | 0 | 0 | 0 | 0 | 0 | 0 | 0 | 1 | 334 | 0.000 |
| | Wide | 2019 | 0 | 0 | 0 | 0 | 0 | 0 | 0 | 0 | 0 | 105 | 2496 | 0.000 |
| SNG | Narrow | 2018 | 0 | 0 | 0 | 0 | 0 | 0 | 1 | 0.01 | 0.04 | 24 | 25 | 0.083 |
| | | 2019 | 0 | 0 | 0 | 0 | 0 | 0 | 1 | 0 | 0 | 228 | 126 | 0.515 |
| | Wide | 2019 | 0 | 0 | 0 | 0 | 0 | 0 | 0 | 0 | 0 | 21 | 122.4 | 0.588 |
| SPB | Narrow | 2017 | 0 | 0 | 0 | 0 | 0 | 0 | 0 | 0 | 0 | 25 | 117 | 0.000 |
| SRS | Narrow | 2017 | 0 | 0 | 0 | 0 | 0 | 0 | 0 | 0 | 0 | 36 | 70 | 0.000 |
| SSC | Narrow | 2017 | 0 | 0 | 0 | 0 | 0 | 0 | 0 | 0 | 0 | 28 | 81 | 0.000 |
| | | 2018 | 0 | 0 | 0 | 0 | 0 | 0 | 0 | 0 | 0 | 24 | 25 | 0.000 |
| SYH | Narrow | 2018 | 0 | 0 | 0 | 0 | 0 | 0 | 0 | 0 | 0 | 55 | 21 | 0.320 |
| | | 2019 | 0 | 0 | 0 | 0 | 0 | 0 | 0 | 0 | 0 | 1 | 86 | 0.000 |
| | Wide | 2019 | 0 | 0 | 0 | 0 | 0 | 0 | 0 | 0 | 0 | 0 | 584.8 | NA |
